# Salvage NAD^+^ Biosynthetic Pathway Enzymes Moonlight as Molecular Chaperones to Protect Against Proteotoxicity

**DOI:** 10.1101/2020.10.27.357780

**Authors:** Meredith Pinkerton, Andrea Ruetenik, Viktoriia Bazylianska, Eva Nyvltova, Antoni Barrientos

**Author notes:** Correspondence to: Antoni Barrientos, Ph.D., Department of Neurology, University of Miami Miller School of Medicine 1420NW, 9^th^ Ave, NRB # 103 Miami. FL-33136. **Classifications** Biological Sciences Neuroscience. **Significance statement** Huntington’s disease and Parkinson’s disease are age-associated neurodegenerative disorders characterized by the misfolding and oligomerization of polyglutamine domains or α-synuclein, leading to neuronal death. To identify cellular protective mechanisms, we have used yeast models expressing the human proteins that recapitulate the basic proteotoxicity principles seen in neurons. Here, we describe four enzymes in the salvage NAD^+^ biosynthetic pathway whose overexpression attenuates proteotoxicity. We show that, under proteotoxic stress, Nma1, Npt1, Pnc1, and Qns1 are recruited as molecular chaperones to prevent the misfolding and support the refolding of polyglutamine domains and α-synuclein, independently of their NAD^+^ synthesis role. Our study connects NAD^+^ metabolism and proteostasis and identifies salvage NAD^+^ biosynthetic proteins as therapeutic targets for neurodegenerative proteotoxicities.

## Abstract

Human neurodegenerative proteinopathies are disorders associated with abnormal protein depositions in brain neurons. They include polyglutamine (polyQ) conditions such as Huntington’s disease (HD) and α-synucleinopathies such as Parkinson’s disease (PD). Overexpression of NMNAT/Nma1, an enzyme in the NAD^+^ biosynthetic salvage pathway, acts as an efficient suppressor of proteotoxicities in yeast, fly, and mouse models. Screens in yeast models of HD and PD allowed us to identify three additional enzymes of the same pathway that achieve similar protection against proteotoxic stress: Npt1, Pnc1, and Qns1. Here, we report that their ability to maintain proteostasis is independent of their catalytic activity and does not require cellular protein quality control systems such as the proteasome or autophagy. Furthermore, we show that, under proteotoxic stress, the four proteins are recruited as molecular chaperones with holdase and foldase activities. The NAD^+^ salvage proteins act by preventing misfolding and, together with the Hsp90 chaperone, promoting the refolding of extended polyQ domains or α-synuclein. We conclude that the entire salvage NAD^+^ biosynthetic pathway links NAD^+^ metabolism and proteostasis and emerges as a target for therapeutics to combat age-associated neurodegenerative proteotoxicities. Our observations also illustrate the existence of an evolutionarily conserved strategy of repurposing or moonlighting housekeeping enzymes under stress conditions to maintain proteostasis.

## INTRODUCTION

Metabolic, mitochondrial, and protein folding abnormalities are a common feature of neurodegeneration and aging. Despite substantial recent progress in understanding the molecular and cellular perturbations associated with these processes, many fundamental questions remain open. As a consequence, no current therapies target the underlying cellular pathologies of age-related neurodegenerative diseases.

Searching for suppressors of proteotoxicity in model organisms has proven valuable for understanding the pathways involved and their contribution to cellular homeostasis and death. The different suppression mechanisms reported to date can be classified into two categories: (i) those suppressing the harmful effects of misfolded/oligomerized proteins upon specific cellular pathways, and (ii) those avoiding the accumulation of misfolded proteins. In the first category, suppression of proteotoxicity is achieved by protecting the cells against some downstream events, such as cytoskeletal instability (1), impaired gene transcription (2), and accumulation of kynurenine pathway toxic metabolites (3). We and others have shown that protection is also achieved by stimulating mitochondrial biogenesis, integrity, and function (2, 4). Specifically, we reported that boosting mitochondrial biogenesis suppresses proteotoxicity in yeast and fly models of Huntington’s disease (HD) (4, 5). In the second category, proteotoxicity can be reduced by enhancing the clearance pathways for the removal of cytoplasmic aggregate-prone proteins through increasing autophagy (6), activating the ubiquitin-proteasome system (7) or the endoplasmic reticulum-associated degradation pathway (8). Toxicity is also ameliorated by modulating the chaperone systems involved in protein refolding, aggregation, and disaggregation (9-15).

In addition to dedicated heat-shock proteins, an unexpected inclusion into the class of molecular chaperones is NMNAT or nicotinamide mononucleotide (NMN) adenylyltransferease. This conserved enzyme is rate-limiting in the human salvage NAD^+^ biosynthetic pathway, in which it reversibly catalyzes the synthesis of NAD^+^ from ATP and NMN. Three NMNAT proteins have been identified in mammals with reported distinct tissue and cellular distribution: NMNAT-1 in the nucleus, NMNAT-2 in the cytoplasm, and NMNAT-3 in the mitochondria (16). NMNAT is essential for maintaining physiological neuronal integrity, and when overexpressed, protects against several neurodegenerative conditions in fly and mouse models, including Wallerian degeneration (17, 18), exotoxicity (19), and neurodegeneration induced by spinocerebellar ataxia 1 (18, 20), Tau (21, 22), or huntingtin (5, 23). It has been hypothesized that increasing NAD^+^ concentration via NMNAT and the subsequent activation of the histone deacetylase SIRT1 plays a role in the protective mechanism (24, 25). It remains controversial whether the neuroprotective effect of NMNAT is mediated by increased NAD^+^ levels or a non-catalytic role of NMNAT (18, 26). However, recent evidence indicates that it could be a contribution of both. It is known that NAD^+^ levels decline with age in mice and humans (27, 28), and treatment with NAD^+^ precursors can significantly improve health by supporting both metabolism and signaling pathways to NAD^+^ sensors such as sirtuin deacetylases and poly-ADP-ribose polymerases (PARPs) (29, 30). However, recent studies in mammalian cellular models of tauopathies (which includes Alzheimer’s disease (AD)) have shown that NMNAT can act as a co-chaperone of the heat shock protein 90 (HSP90) to mediate proteostasis, independently of NAD^+^ levels (31). Moreover, structural approaches have revealed that NMNAT uses its enzymatic pocket to specifically bind the phosphorylated sites of pTau, in competition with the enzymatic substrates of NMNAT (32). Furthermore, the capacity of NMNAT to refold misfolded proteins requires a unique C-terminal ATP site, activated in the presence of HSP90 (31).

We and others have screened for suppressors of proteotoxicity using yeast models of polyglutamine (polyQ) disorders such as HD and α-synucleinopathies such as Parkinson’s disease (PD). These models recapitulate the critical events preceding cell death that manifest in human conditions, such as protein misfolding and aggregation and mitochondrial dysfunction (4, 5, 8, 33-35). Using these models, we have demonstrated that the yeast homologs of NMNAT, Nma1, and Nma2, act as powerful suppressors of proteotoxicities (35). Furthermore, we revealed three other enzymes in the yeast salvage NAD^+^ biosynthetic pathway with a similar capacity (35). These are Npt1 (nicotinate-nicotinamide phosphoribosyltransferase), Pnc1 (nicotinamidase), and Qns1 (NAD^+^ synthase). We reported that the suppression mechanism is independent of NAD^+^ levels or sirtuins and does not necessarily require mitochondrial oxidative phosphorylation (5, 35).

Here, we present data that strongly support the hypothesis that salvage NAD^+^ biosynthetic proteins Nma1/2, Npt1, Pnc1, and Qns1 have intrinsic chaperone activity, independent of their catalytic function. We demonstrate that these proteins have holdase and foldase capacity, which rely on different domains of the proteins. Deleting the C-terminal domain of Nma1 markedly attenuates the Nma1 capacity to suppress 103Q- or α-synuclein-induced proteotoxicity. Furthermore, deleting the HSP90 chaperones Hsc82/Hsp82 abrogates the capacity of Nma1/2, Npt1, Pnc1, or Qns1 to protect against 103Q, although it does not limit their ability to attenuate α-synuclein-induced toxicity. Thus, salvage NAD^+^ biosynthetic proteins and HSP90 chaperones may cooperate to maintain proteostasis in a substrate-specific manner. In yeast, the mechanism by which Nma1/2, Npt1, Pnc1, or Qns1 clear misfolded/oligomerized proteins is independent of proteasomal degradation and autophagy. *In cellulo* and *in vitro* chaperone assays indicate that the four salvage NAD^+^ biosynthetic proteins can perform holdase and foldase activities. These observations indicate that the salvage NAD^+^ biosynthetic proteins act as chaperones to prevent misfolding and assist in refolding misfolded polyQ domains and α-synuclein. Our studies identify new therapeutic targets to treat patients suffering from neurodegenerative proteotoxicities such as HD and PD.

## RESULTS AND DISCUSSION

### Catalytically inactive forms of NAD^+^ salvage pathway proteins retain protective activity against 103Q and α-synuclein toxicity in yeast

We and others have previously described that heterologous expression in the yeast *Saccharomyces cerevisiae* of either human mutant PolyQ domains (103Q) or α-synuclein fused to green fluorescent protein (GFP) under the control of a *GAL1* promoter induced by growth in media containing galactose (YPGAL) results in significant growth defects (5, 33-36). We have also reported that in yeast models of HD and PD, overexpression of hemagglutinin (HA)-tagged versions of the *NMNAT* yeast homologs, *NMA1 and NMA2*, or other genes in the NAD^+^ biosynthetic salvage pathway, including *QNS1*, *NPT1,* and *PNC1* (**Fig 1A and 1B**) suppresses 103Q- and α-Syn-induced cytotoxicity, through a mechanism that involves extensive clearance of non-native proteins, without affecting the expression levels of mutant PolyQ or α-synuclein, as reported (35). Importantly, we demonstrated that the suppression mechanism does not require the presence of a functional NAD^+^ biosynthesis salvage pathway because the deletion of one of the enzymes (*NPT1*), which lowered cellular NAD^+^ levels by ~2.2 fold, did not affect the suppression of 103Q-induced toxicity by NMA1/2 or other components of the pathway (35).

**Fig 1.**
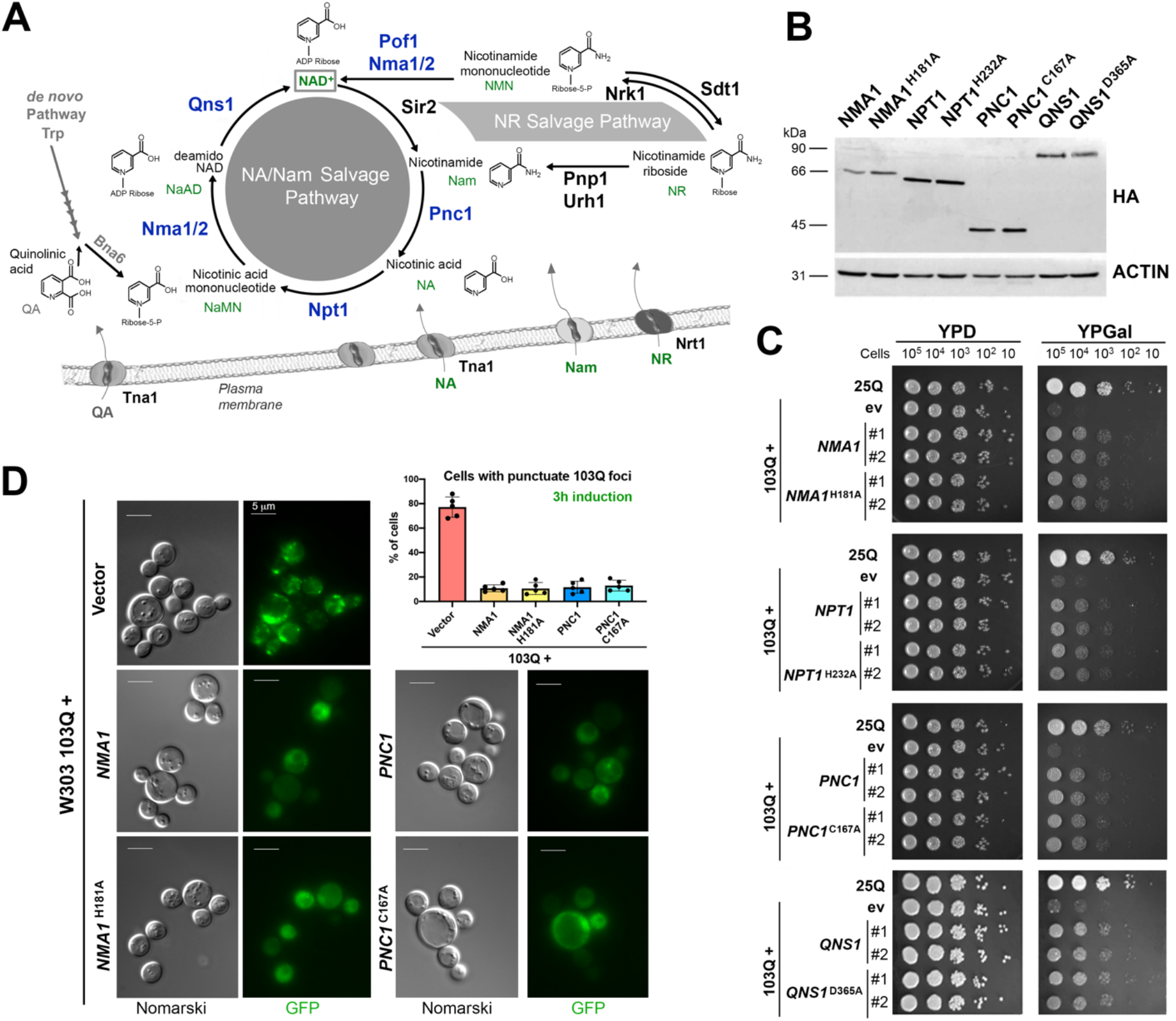
Catalytically inactive NAD^+^ salvage enzymes suppress 103Q-induced proteotoxicity in yeast. (**A**) Overview of the NAD^+^ salvage pathway in yeast. (**B**) Immunoblot analysis of expression of catalytically active and inactive HA-tagged NAD^+^ salvage proteins. ACTIN was using as a loading control. (**C**) Serial dilutions growth test of wild-type cells expressing non-toxic 25Q or toxic 103Q, and 103Q cells expressing WT or catalytically inactive variants of the indicated salvage NAD^+^ pathway genes, under the control of a galactose inducible promoter, in solid non-inducing media (YPD) and inducing galactose media (YPGal). #1 and #2 are two independent clones. Pictures were taken after 2 days of growth. (**D**) Visualization of PolyQ-GFP fusion protein expression and aggregation. The indicated strains were induced for 3 h in complete medium with 2% galactose, mounted on slides and visualized by fluorescence microscopy. Bar is 5 μm. The graph quantifies the cells with at least one 103Q aggregation focus. Three hundred cells were counted in each case. Error bars are the mean ± SD, of five independent assessments with *P*-values from comparisons to 103Q + vector strain denoted by \**P* < 0.01.

To further discern the properties of these salvage NAD^+^ biosynthetic proteins required to confer protection against proteotoxicity, we mutated the catalytic core of each protein by targeting previously identified relevant residues. The yeast Nma1-H181A (human H24A) (26, 37), Npt1-H232A (human H247A) (38), and Pnc1-C167A mutations affect the catalytic center of each enzyme and eliminate their function (39). Qns1 has two functional domains; mutants in the glutaminase domain (C175A) or the synthetase allele D365A are non-functional *in vivo* (40). Catalytic mutants retain less than 5% of WT activity as reported (26, 37-40) and as measured in purified recombinant proteins (see below). When the proteins were expressed in the HD and PD yeast models, the catalytically active and inactive variants of each protein accumulated at similar levels (**Fig 1B**). No difference was observed between the capacity of the catalytically active and inactive variants to suppress the growth defect induced by 103Q (**Fig 1C**) or α-Syn (**supplemental Fig S1**).

Catalytically active and inactive variants showed a similar ability to promote the clearance of non-native proteins (**Fig 1D**and **2**). Using fluorescence microscopy, we followed GFP-tagged mutant PolyQ expression and oligomerization in strains overexpressing wild-type or mutant variants of *NMA1*, *NMA2*, *PNC1, QNS1,* or carrying an empty vector (YEp352). After 1 h of protein expression induction in galactose medium, the 103Q-GFP signal was primarily diffuse in the cytoplasm, forming a few small aggregates. The number and size of the aggregates progressed with time, and after only 3 h of induction, the protein formed large aggregates (**Fig 1D**). Overexpression of catalytically active or inactive variants of *NMA1*, *NPT1, PNC, and QNS1* was able to attenuate the fluorescence signal of both soluble and aggregated mutant PolyQ as well as the number of aggregates (**Fig 1D**). Approximately 75% of the 103Q cells contained at least one 103Q-GFP punctuate focus after 3 h of 103Q-GFP expression. This value was reduced to ∼10–15% in cells overexpressing wild-type or mutant *NMA1* or *PNC1* (**Fig 1D**). Overexpression of *NMA1*, *NPT1*, *PNC1, or QNS1* did not affect the level of expression of 103Q, as we have previously reported (35), suggesting that they prevent 103Q misfolding or increase its degradation. To assess the effect of catalytically inactive NAD^+^ proteins on the kinetics of 103Q oligomerization, we performed immunoblot analysis of protein extracts from cells expressing 103Q alone or in combination with the suppressor genes (**Fig 2A**). In agreement with our fluorescence microscope analyses, we observed a time-dependent protein aggregation pattern, as previously reported (33, 35). 103Q strain, after 2 h of expression, 103Q-GFP proteins were found both at their expected size and forming SDS-insoluble oligomers (**Fig 2B**). At later time points, 103Q-GFP was detected only as part of SDS-insoluble large oligomers trapped in the higher portions of the stacking gel (**Fig 2B**). As controls, 25Q accumulated in a soluble state (**Fig 2B**), and the deletion of *hsp104* largely prevented 103Q oligomerization (**Fig 2C**), as reported (41). Treatment of 103Q cells with the NAD^+^ precursor nicotinamide did not change the pattern of 103Q oligomerization. This result is consistent with the observation that boosting NAD^+^ biosynthesis with NAD^+^ precursors (nicotinamide, nicotinic acid, or nicotinamide riboside) does not ameliorate either 103Q- or α-Syn-induced proteotoxicity, or the protection exerted by overexpression of salvage NAD^+^ biosynthetic proteins (**supplemental Fig S2**). The 103Q oligomerization pattern changed dramatically when catalytically active or inactive forms of NAD^+^ biosynthetic proteins were overexpressed in 103Q cells. In these cell lines, we detected lower amounts of SDS-insoluble large oligomers at several time points, accompanied by increased signal corresponding to either degradation products or intermediate states of protein oligomerization (**Fig 2E-H**). Note that the overexpressed proteins, which are tagged with ZZ, a Protein A fragment, are also detected in these immunoblots.

**Fig 2.**
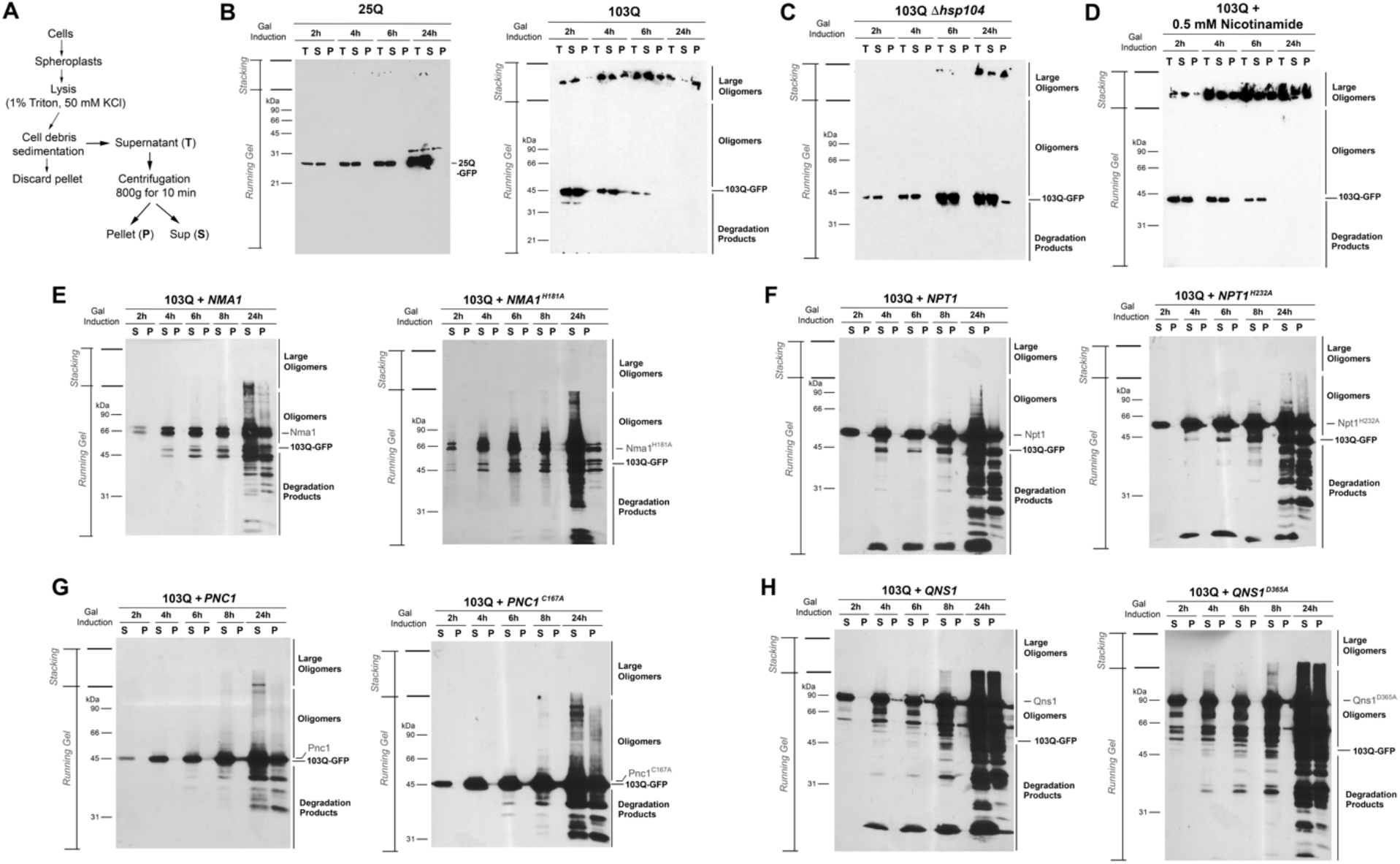
Suppression by catalytically active or inactive NAD^+^ biosynthetic proteins equally involves the clearance of cytotoxic protein species. (**A**) Scheme of the protocol followed to isolate aggregates from cultures expressing the short PolyQ and the long PolyQ constructs. Samples corresponding to the cell lysates after removal of the cell debris sedimented by gravity (T), the pellet (P) and supernatant (S) fractions resulting from centrifugation at 800 *g* for 10 minutes were taken and kept frozen until further use. (**B-H**) Detection of PolyQ-GFP fusion proteins. Wild-type W303-1A cells carrying integrative plasmids expressing 25Q-GFP or 103Q-GFP fusion proteins under the control of a *GAL1* promoter were grown on minimal synthetic media supplemented with prototrophic requirements and 2% galactose to induce protein expression. After several times of induction, the cultures were processed as described in (A). PolyQ-GFP fusion proteins were detected by western blot using equivalent amounts of the T, S and P fractions. After separation on a polyacrylamide gel, proteins were transferred to nitrocellulose paper. The fusion proteins were detected using an anti-GFP monoclonal antibody (Clontech, Palo Alto, CA) and visualized with anti-mouse IgG peroxidase-conjugated secondary antibody (Sigma) using the Super Signal chemiluminescent substrate kit (Pierce).

The data presented in this section allowed us to conclude that the ability of NAD^+^ salvage proteins to protect against proteotoxicity in yeast is independent of their catalytic activity.

### NAD^+^ salvage proteins function as chaperones with both holdase and foldase activities

To assess if NAD^+^ salvage proteins can directly bind to substrates and reduce protein aggregation in the absence of other proteins, we performed a cell-free chaperone activity assay based on the fact that, under thermal stress, citrate synthase (CS) aggregates leading to increased optical absorbance (31). In this assay, the use of purified candidate chaperone proteins allows assessing whether they can exert holdase activity to maintain CS in thermostable conformation in the absence of other chaperones, co-chaperones, or ATP. For this purpose, we expressed in *E. coli* and purified 6xHis-tagged recombinant catalytically active and inactive Nma1, Npt1, Pnc1, and Qns1 proteins (**Fig 3A**). The four wild-type yeast proteins were previously expressed in *E. coli*, purified, and found to be catalytically active (38, 40, 42-44). We have measured the activity of wild-type and mutant Nma1 and Pnc1 using spectrophotometric coupled enzyme assays (39, 45) and demonstrated that the mutant proteins are inactive. Their measured activities were either undetected or lower than 1% of wild-type. In the CS-aggregation assay, all of the proteins, catalytically active or inactive, prevented thermally induced CS aggregation in a dose-dependent manner (**Fig 3B**). As a negative control, we used lysozyme, which did not have any chaperone activity (**Fig 3B**).

**Fig 3.**
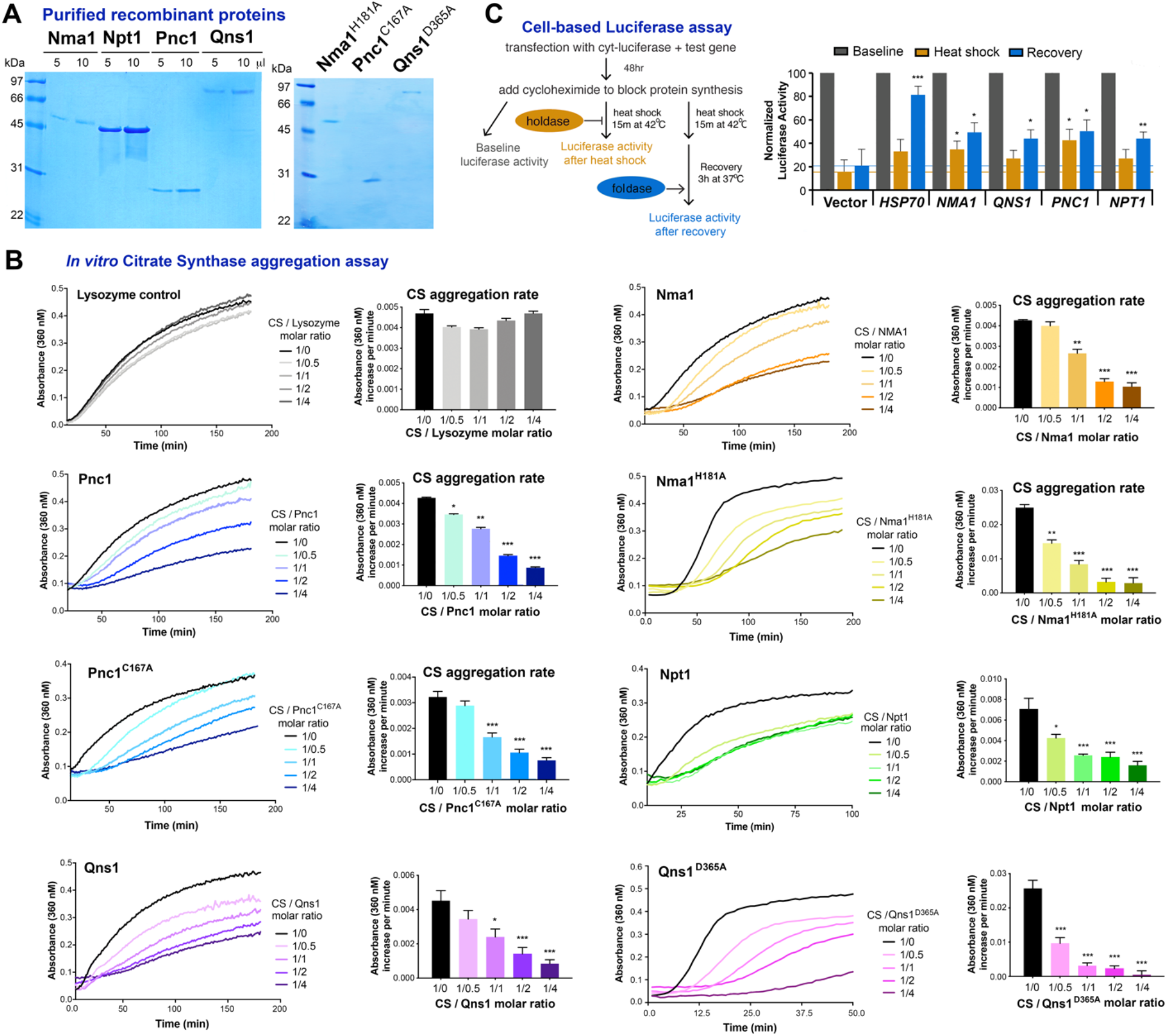
NAD^+^ salvage proteins function as chaperones with both holdase and foldase activities. (**A**) Coomassie blue stained gel showing the recombinant proteins purified as 6xHis fusion proteins from *E. coli*. (**B**) Citrate synthase (CS) aggregation assay for NMA1, NPT1, PNC1, and QNS1 WT and catalytically inactive mutants). At least five independent repetitions were performed for each protein. The bar graphs represent the average aggregation rates of citrate synthase alone (1μM citrate synthase) or 1μM citrate synthase plus the indicated amount of protein of interest during the period in which the rate of citrate synthase aggregation is linear. Error bars are the mean ± SEM. *p ≤ 0.01, ** p ≤ 0.001, ***p ≤ 0.0001. (**C**) The diagram illustrates simplified experimental procedure of cell-based luciferase denaturation and refolding assay. The graph summarizes the chaperone activity of HSP70, and salvage NAD^+^ biosynthetic proteins NMA1, NPT1, PNC1, and QNS1. Grey bars show baseline luciferase activity. Orange bars show luciferase activity immediately after heat shock, while blue bars show activity after recovery. Error bars are the mean ± SD, of three independent assays with *P*-values from comparisons to Vector cell line denoted by \**P* < 0.05, ** p ≤ 0.01, ***p ≤ 0.001.

To further assess the chaperone activity of yeast NAD^+^ salvage proteins, we conducted a cell-based luciferase assay (31), which determines the ability of candidate proteins in preventing luciferase misfolding or facilitating luciferase refolding in intact cells with cellular protein repair machinery. In this assay, treatment with cycloheximide to block protein synthesis, followed by heat denaturation (at 42°C for 15 min), renders the endogenous chaperone machinery incapable of preventing luciferase aggregation and refolding (**Fig 3C**). This allows for direct measurement of the chaperone activity of the introduced test protein by its ability to prevent luciferase from undergoing heat shock-induced denaturation (holdase activity) and to promote its proper refolding (foldase activity) during recovery (at 37°C for 3 h). We found that the four NAD^+^ salvage proteins have holdase activity comparable or higher to that of HSP70 (**Fig 3C**), and robust foldase activity approximately 60% to HSP70 (**Fig 3C**).

### The proteasome, autophagy, or yeast chaperones proteins Hsp26, Hsp42, or Hsp70 (Ssa1) are not required for protection by NAD^+^ salvage pathway proteins

The ability of the yeast salvage NAD^+^ biosynthetic proteins to protect against cytotoxicity could be intrinsic or depend on other protein quality control cellular systems (35). In *Drosophila,* the Nma1/2 homolog NMNAT directly interacts with phosphorylated Tau and promotes the ubiquitination and clearance of toxic Tau species in the nervous system in a tauopathy model (21). Moreover, *Drosophila* NMNAT significantly mitigates mutant huntingtin (138Q)-induced neurodegeneration by reducing mutant huntingtin aggregation through promoting autophagic clearance (23). To reveal the potential protective pathways involved in the NAD^+^ biosynthetic protein-mediated clearance of mutant polyQ and α-Syn oligomers in yeast, we tested pathways that are affected by and mediate proteotoxic stress, the unfolded protein response (UPR) or enhance autophagy, (8, 46-48). We observed that protection could not be abolished or diminished by inhibiting the proteasome with 10μM or 50μM MG132 (**supplemental Fig S3**), indicating that the proteasome is not required for protection by NAD^+^ salvage pathway proteins. We further established that blocking autophagy in yeast by introducing a Δatg32 mutation or blocking mitophagy by introducing a Δatg12 mutation did not modify the efficacy of proteotoxicity suppression by NAD^+^ salvage pathway proteins (**supplemental Fig S4** and **S5**).

The salvage NAD^+^ biosynthetic proteins could also protect by enhancing the function of other chaperones (e.g., small Hsps or Hsp70 families) known to modify the folding, aggregation, and toxicity of mutant polyQ in yeast (10, 49-52). Hsp26 and Hsp42 are known to sequester misfolding proteins in near-native conformation for cellular protection and efficient refolding (53) and to cooperate with Hsp140 and the Hsp70 chaperone Ssa1 in protein disaggregation (10). However, the absence of Hsp26, Hsp42 or Ssa1 did not affect the ability of salvage NAD^+^ biosynthetic proteins to protect against proteotoxicity (**supplemental Fig S6** and **S7**). The lack of effect of Ssa1 could be explained by the fact that the major cytosolic Hsp70 family in yeast, the Hsp70-Ssa, consists of four members of Ssa (constitutively expressed Ssa1-2 and stress-induced Ssa3-4), which are functionally redundant to some degree as the expression of at least one family member is essential for growth (54).

These data indicate that the NAD^+^ proteins may confer protection through a pathway that does not require Hsp26, Hsp42, or Ssa1. Alternatively, when overexpressed, the protective proteins may substitute for the roles that Hsp26, Hsp42, or Ssa1 usually play.

### HSP90-family chaperones are essential for the protective activity of salvage NAD^+^ proteins against polyQ-induced proteotoxicity

In the yeast *S. cerevisiae*, there are two HSP90 isoforms, encoded by the genes *HSC82* and *HSP82* (55). The deletion of Hsc82 in the 103Q background did not prevent protection by NAD^+^ proteins but slightly attenuated their efficacy (**supplemental Fig S8**). The double deletion Δ*hsp82*Δ*hsc82* did not affect cell survival in YPD media. Still, it enhanced 103Q toxicity in inducing YPGal media and completely abolished the capacity of salvage NAD^+^ biosynthetic proteins to suppress proteotoxicity (**Fig 4**). These results agree with a recent study in mammalian cellular models of tauopathies showing that NMNAT2 can act as an HSP90 co-chaperone to mediate proteostasis, independently of NAD^+^ levels (31). Unexpectedly, however, the double deletion Δ*hsp82*Δ*hsc82* did not interfere with the suppression of α-Syn-induced toxicity, suggesting that the HSP90 co-chaperone activity of salvage NAD^+^ proteins does not operate for all substrates.

**Fig 4.**
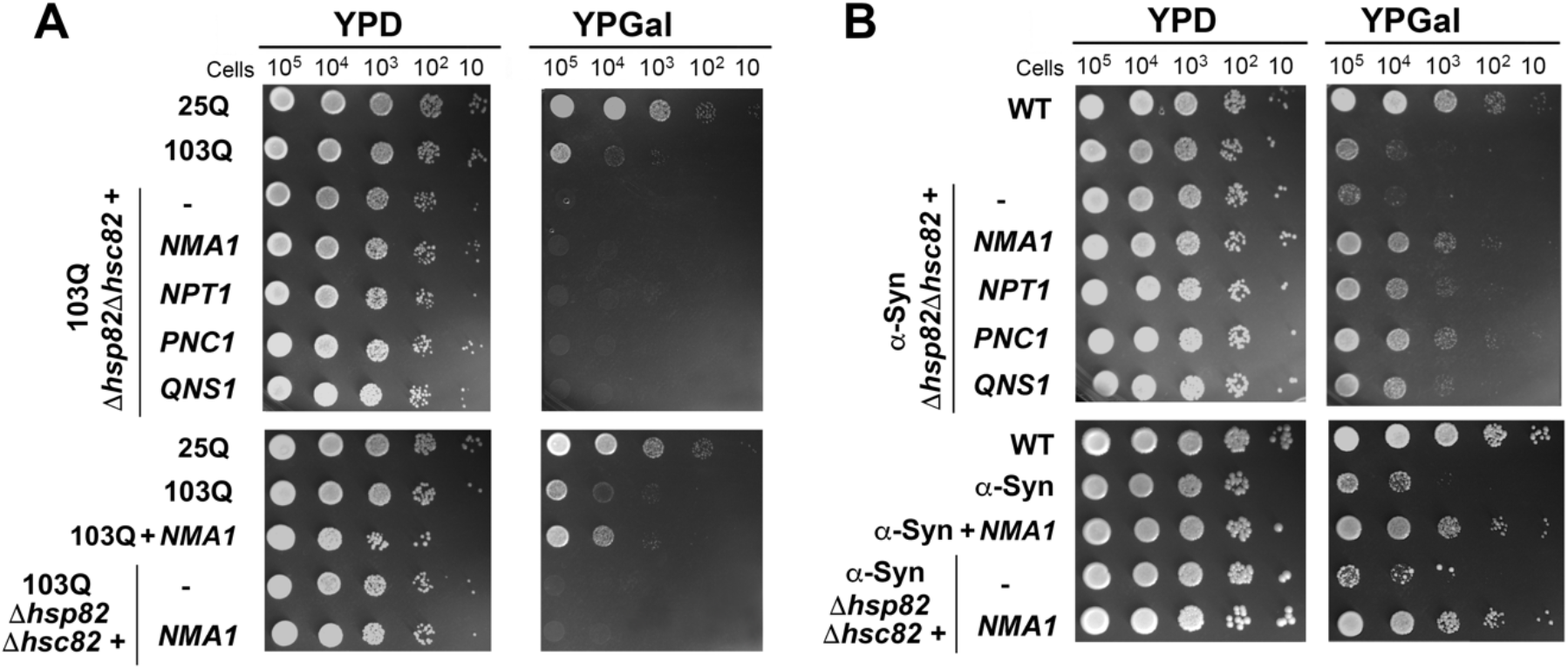
Double deletion of the two HSP90 chaperones, Hsp82 and Hsc82, prevents the suppression of 103Q-induced but not α-Syn-induced toxicity by NAD^+^ salvage proteins. **(A)** Serial dilutions growth test of wild-type cells expressing non-toxic 25Q or toxic 103Q, and 103Q cells deleted or not for *hsp82* and *hsc82*, expressing the indicated salvage NAD^+^ pathway genes, under the control of a galactose inducible promoter, in solid non-inducing media (YPD) and inducing galactose media (YPGal). Pictures were taken after 2 days of growth. **(B)** Same as in (A) but for α-Syn-expressing strains.

### The C-terminus of yeast protein Nma1 is essential for protective activity

To elucidate whether the catalytic and chaperone functions of NAD^+^ salvage pathway proteins occur in distinct domains of the protein is a very relevant question. We have shown that mutations in the catalytic core of each enzyme block their activity but not their capacity to act as molecular chaperones and protect against proteotoxicity (**Fig 1-3**). However, structural approaches have revealed that human NMNAT2 uses its enzymatic pocket to specifically bind the phosphorylated sites of pTau, in competition with the enzymatic substrates of NMNAT (32). Here, we have focused on Nma1 to perform structure-function relationship studies to identify potential domains required for its chaperone activity while not affecting its catalytic activity. The amino acids we chose to mutate were selected based on human NMNATs studies (56-58) and resulted in four Nma1 variants that we called H3, I-H, NI, and Δct (**Fig 5A and supplemental Fig S9**). Nma1/NMNAT has a structural similarity with *E. coli* chaperones UspA and Hsp100 (56). NMNAT α-Helix 3 (H3) has the best overlapping score and includes Nma1 amino acids 252-TAKVLDH-259 (56). In the Nma1 H3 mutant, the conserved amino acids VL were changed to EE. Nma1 α-Helix 4 contains amino acids 220-RTGSDVRSFL-229 and occupies an internal position in the hexameric NMNAT1 structure (I-H). In the Nma1 I-H mutant, the sequence was changed to 220-<u>A</u>TGS<u>A</u>VRSF<u>A</u>-229. In the Nma1 NI mutant, two conserved residues, Nma1 341-NI-342, located in a hotspot region for human NMNAT1 mutations associated with Leber congenital amaurosis 9 (LCA9) despite not affecting catalytic activity (57), were changed to −AA. Finally, because the capacity of NMNAT2 to refold misfolded proteins requires its C-terminal domain (31), we created a Nma1Δct variant missing aa 375-394 from the Nma1 401aa wild-type protein.

**Fig 5.**
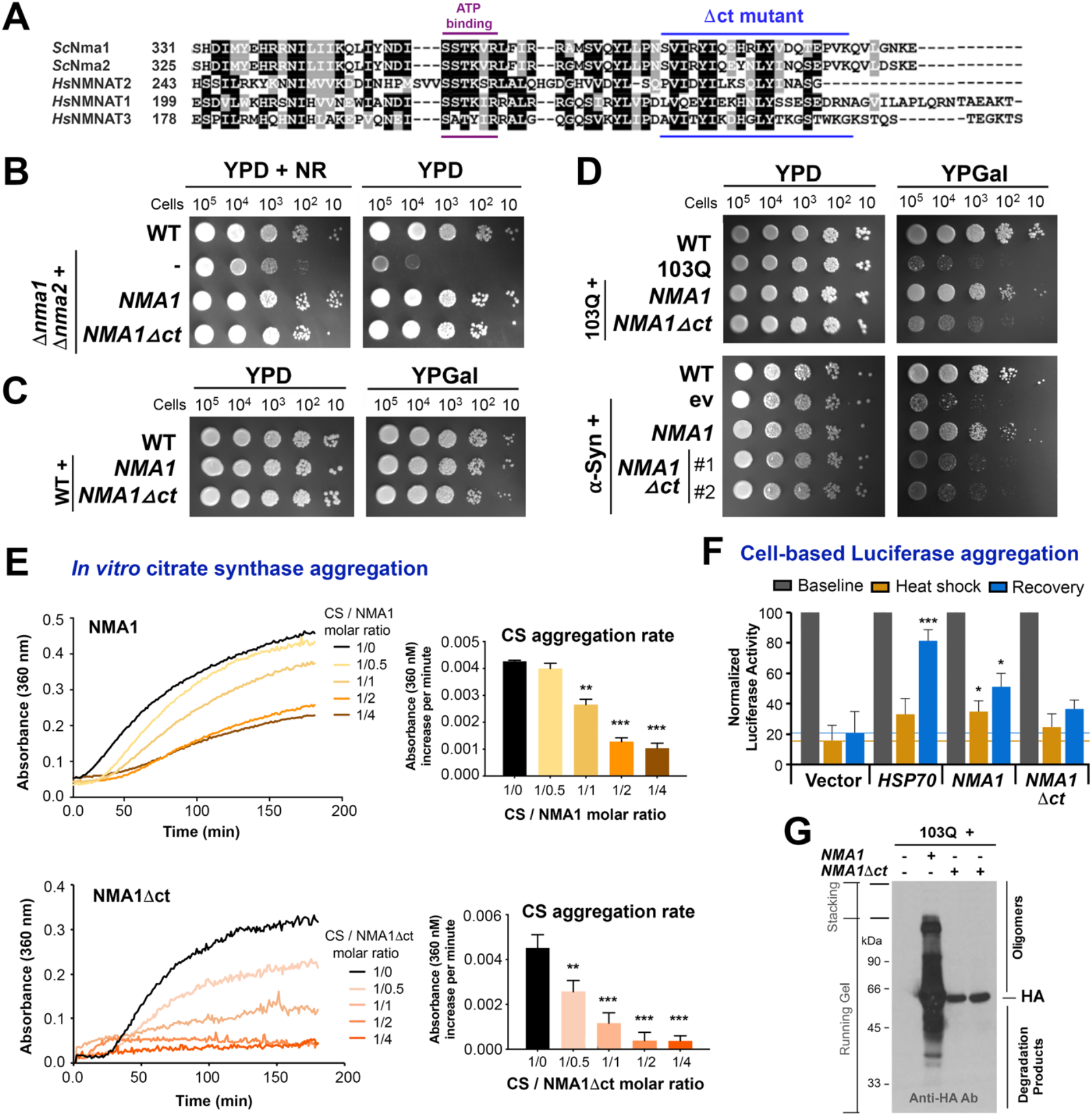
The C-terminus of Nma1 is required for full protection against proteotoxicity. (**A**) Multiple protein alignment of *Saccharomyces cerevisiae* Nma1 and Nma2 and *Homo sapiens* NMNAT1, NMNAT2, and NMNAT3 using the Clustal Omega program from EMBL-EBI. The C-terminal fragment deleted in the *NMA1Δ*ct yeast strain is marked in blue. (**B-D**) Serial dilutions growth tests as in Figure 4 to assess: (**B**) the ability of Nma1Δct to restore growth of a double mutant Δnma1Δnma2 in YPD medium without nicotinamide riboside (NR); (**C**) the effect of *NMA1* or *NMA1Δct* overexpression in wild-type (WT) W303 cells; (**D**) the ability of Nma1Δct to protect against 103Q-induced proteotoxicity. (**E**) Citrate synthase (CS) aggregation assay for recombinant Nma1 and Nma1Δct performed as in Fig 3B. (**F**) Cell-based luciferase denaturation and refolding assay performed as in Fig 3C. The graph summarizes the chaperone activity of HSP70, NMA1 and Nma1Δct. Grey bars show baseline luciferase activity. Orange bars show luciferase activity immediately after heat shock, while blue bars show activity after recovery. Error bars are the mean ± SD, of three independent assays with *P*-values from comparisons to Vector cell line denoted by \**P* < 0.05, ** p ≤ 0.01, ***p ≤ 0.001. (**G**) Cellular extracts from the indicated strains after 6h of induction in YPGal medium were prepared as reported (33), and analyzed by immunoblotting using an anti-HA Ab to detect WT and Nma1Δct.

Yeast expressing each of these mutated Nma1 variants were then screened for their ability to protect against 103Q cytotoxicity. The H3, I-H, and NI Nma1 mutants were found protective, albeit that the efficacy of the H3 variant was slightly attenuated (**supplemental Fig S9**). On the contrary, protection against both 103Q and α-Syn-induced proteotoxicity by the Nma1Δct variant was largely precluded (**Fig 5D**). The Nma1Δct protein retains catalytic activity as it was able to restore the growth of a double mutant Δnma1Δnma2 strain in media lacking exogenous nicotinamide riboside (NR) (**Fig 5B**), and its overexpression in wild-type cells does not induce any gross dominant-negative effect in cell growth (**Fig 5C**). Purified recombinant Nma1Δct (**supplemental Fig S10**) prevented heat-induced CS aggregation *in vitro*, suggesting retention of its holdase activity (**Fig 5E**). This was confirmed by the *in cellulo* luciferase assay, which showed that Nma1Δct have holdase activity 80% of Nma1 (**Fig 5F**), but poorer foldase activity approximately 60% to Nma1 (**Fig 5F**). In 103Q yeast cells, we further confirmed that the Nma1Δct protein was successfully and stably expressed (**Fig 5G**). Notably, upon 6h of 103Q induction in galactose medium, Nma1 showed a pattern of oligomerization and degradation similar to 103Q (**Fig 2E**), suggesting an interaction. On the contrary, Nma1-Δct was stably expressed and did not follow that pattern (**Fig 5G**), which could result from a failure to stably bind 103Q.

The attenuation of the protection capacity by Nma1Δct protein results from a loss of its chaperone capacity, particularly its foldase activity. In *Drosophila* NMNAT (18) or human NMNAT2 (31), the deletion of the entire C-terminus of the proteins (aa 200-308) suppresses their foldase activity. A selective loss of foldase activity by the Nma1Δct protein could be related to the nearby presence of a highly conserved ATP binding site 354-SSTKVR-359 in Nma1 (31)(**Fig 5A**), whose folding could be altered. ATP binding is known to enhance the chaperone activity of the large heat shock proteins by suppressing their unfolding and oligomerization (59, 60). The loss of foldase activity has also been observed in NMNAT2 by deleting only the ATP binding motif (aa 269-SSTKVR-274), which also precludes the recruitment of HSP90 chaperones (31). In yeast, Nma1, Npt1, Pnc1, and Qns1 also have ATP- or phosphate-binding motifs in or near their C-terminus, which could be the commons feature among these proteins to explain their foldase activity. However, deletion of their ATP binding motif did slightly attenuate the capacity of Nma1 and Qns1 to maintain proteostasis (**Fig S11**), indicating the complete C-terminal domain, including the ATP binding site, is crucial to refold of misfolding proteins.

### CONCLUSION

Accumulation of misfolded, oligomerized, and aggregated proteins is a hallmark of age-associated neurodegenerative disorders in humans. Several proteins of unrelated sequences, normally soluble, such as polyglutamine (polyQ) expanded huntingtin in HD, α-Syn in PD, and Tau and Aβ in AD, form neurotoxic amyloids that are morphologically similar and have a core structure of characteristic cross β-conformation (61-66). As the formation of protein oligomers and aggregates involve conformational changes in proteins, various chaperones have been extensively studied for their role in the process and as therapeutic targets, including the small heat shock proteins Hsp26 and Hsp42, as well as Hsp70, Hsp90, and Hsp100 family of proteins (9, 10, 49, 67-70). In addition to heat shock proteins, the salvage NAD^+^ enzyme NMNAT has been shown to act as a stress-response protein that works as a chaperone for neuronal maintenance and protection in fly and mouse models of neurodegenerative disorders (18, 31, 32, 56). Using yeast models of HD and PD, we have previously shown that in addition to the NMNAT homologs Nma1 and Nma2, three additional salvage NAD^+^ biosynthetic proteins, namely Npt1, Pnc1, and Qns1, suppress 103Q and α-Syn-induced proteotoxicity by promoting the clearance of misfolded/oligomerized proteins (35). In this manuscript, we show that (i) the anti-proteotoxicity role of Nma1, Npt1, Pnc1, and Qns1 is independent of their enzymatic activity in the salvage NAD^+^ biosynthetic pathway, (ii) they act as chaperones with holdase and foldase activities, and (iii) they can cooperate with HSP90-family proteins in a misfolding substrate-dependent manner. A model of proteotoxicity suppression by salvage NAD^+^ biosynthetic proteins is presented in **Figure 6.**

In addition to their primary function of preventing protein aggregation, the role of chaperones in directing terminally misfolded proteins to either proteasome or autophagy is well established (71). Our data indicate that the yeast salvage NAD^+^ biosynthetic proteins retain their protection capacity independently of the cellular protein degradation machinery. This contrasts with previous observations regarding the role of NMNAT made in fly models of tauopathy and HD in which the proteasome and autophagy are activated, respectively, when NMNAT is overexpressed (21, 23). Our data suggest that if Nma1, Npt1, Pnc1, and Qns1 were more efficient in preventing misfolding and supporting refolding of 103Q or α-Syn in the yeast than in the fly models (*e.g.,* if the overexpression levels are higher), the cellular protein quality control systems could be differently recruited in the two model organisms.

**Fig 6.**
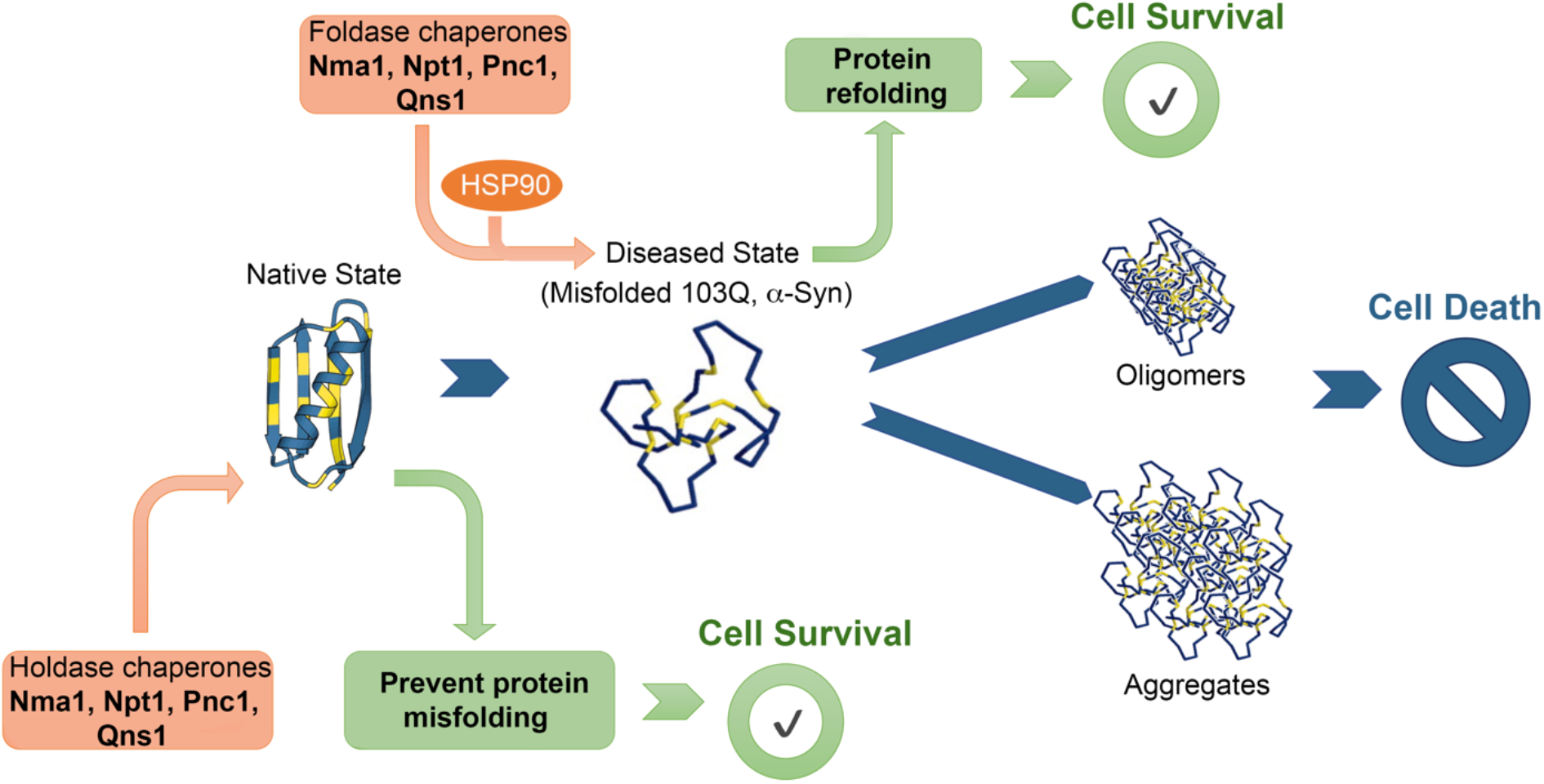
Model of proteotoxicity suppression by salvage NAD^+^ biosynthetic proteins. Under proteotoxic stress, salvage NAD^+^ biosynthetic proteins Nma1, Nma2, Npt1, Pnc1 and Qns1 are recruited to act as molecular chaperones. They display foldase activity to prevent protein misfolding and cooperate with HSP90-type chaperones to display foldase activity and promote protein refolding. In this way, NAD^+^ biosynthetic proteins contribute to minimize formation of toxic oligomers and aggregates to promote cell survival. See further explanation in the text.

The role of NMNAT proteins as neuroprotective chaperones independent of NAD^+^ synthesis has been controversial mainly because studies using models of Wallerian axonal degeneration proposed that NMNAT overexpression maintains axonal integrity by enhancing NAD^+^ synthesis and preventing the accumulation of the precursor NMN (72-74). However, it was later observed that NAD^+^ depletion upon axonal injury resulted from fast consumption by the prodegenerative SARM1 enzyme, which is blocked by NMNAT1 without enhancing NAD^+^ synthesis (75). However, the NAD^+^ biosynthetic function of NMNATs can be relevant in a context-dependent manner. For example, the deletion of NMNAT2 in cortical neurons renders them sensitive to both proteotoxic stress and excitotoxicity triggered by sustained neuronal depolarization (31). Still, whereas the chaperone activity of NMNAT2 is required to prevent proteotoxicity, its enzymatic function protects against excitotoxicity (31).

On the other hand, NAD^+^ levels steadily decline with age in mice and humans (27, 28), resulting in altered metabolism and increased disease susceptibility (76). Restoration of NAD^+^ levels in old or diseased animals can promote health and extend lifespan (76). Treatment with NAD^+^ precursors can significantly improve health by supporting both metabolism and signaling pathways to NAD^+^ sensors such as sirtuin deacetylases and poly-ADP-ribose polymerases (PARPs) (29, 30). Supplementation with NAD^+^ precursors could, in principle, add to the neuroprotective role of salvage NAD^+^ biosynthetic proteins acting as chaperones against PolyQ or α-Syn-induced toxicity. However, in our yeast models, NA, Nam, or NR did not affect polyQ toxicity or suppression by Nma1, Npt1, Pnc1, or Qns1. Whereas the decline of NAD^+^ levels with age is significantly contributed by its degradation by the NADase CD38 (27, 28), it remains to be determined whether it is also contributed by the repurposing of NAD^+^ salvage enzymes as chaperones to combat age-associated disruption of proteostasis.

It also remains to be understood how the four salvage NAD^+^ biosynthetic proteins switch between their two functional modes, catalytic in NAD^+^ synthesis *vs.* cytoprotective chaperones. However, recent data have shed light on the structural determinants of NMNAT moonlighting roles. Structural studies of NMNAT bound to pTau have shown that NMNAT is distinct from canonical molecular chaperones (32). NMNAT adopts its enzymatic pocket to specifically bind the phosphorylated sites of pTau, without inducing large conformational changes and can be competed out with the NMNAT enzymatic substrates (NMN and ATP) (32). Because NMNAT uses the same domain, even a shared pocket, for the binding of ATP and pTau, ATP hydrolysis is not required for the chaperone function of NMNAT. Additionally, NMNAT functions as an Hsp90 co-chaperone for the specific recognition of pTau over Tau (31, 32). Our data indicate that yeast Nma1 might also recruit the HSP90 chaperones Hsc82/Hsp82 via its C-terminal domain since both are essential for the Nma1-driven attenuation of 103Q-induced toxicity.

In conclusion, the entire salvage NAD^+^ biosynthetic pathway links NAD^+^ metabolism and proteostasis and is emerging as a potential critical target for therapeutics to combat age-associated neurodegenerative proteotoxicities.

## MATERIALS AND METHODS

### Yeast strains and media

The *S. cerevisiae* strains used were the wild-type W303-1A (*MATa ade2-1 his3-1,15 leu2-3,112 trp1-1 ura3-1*) and the previously reported isogenic 25Q, 103Q, and α-Syn strains (35), and the previously reported W303-1A strains containing chromosomally integrated *N*-terminus of human huntingtin with 103Q or wild-type α-synuclein fused to GFP expressed under the control of the *Gal1* promoter (33, 35).

The plasmids overexpressing *NMA1*, *NMA2*, *NPT1*, *PNC1*, *QNS1*, and *TNA1* were obtained from Open Biosystems (Thermo Scientific, Lafayette, CO, USA), and transformed into the indicated strains. *ATG12, ATG32, SSA1, HSP26, HSP42, HSP82, and HSC82* were deleted by replacing the open reading frame with a *KANMAX4* cassette. To generate a double deletion mutant Δ*hsc82*Δ*hsp82*, the HSP82 open reading frame was replaced with a HIS5 cassette in the Δ*hsc82* strains. Catalytic mutant variants of *NMA1, NPT1, PNC1*, and *QNS1,* and all mutant forms of NMA1 were created using a Q5 site-directed mutagenesis kit (New England Biolabs, Ispwich, MA, USA).

The following complete YP (1% yeast extract, 2% peptone) and minimum WO (0.67% nitrogen base w/o amino acids) media were used routinely to grow yeast: YPD (+2% glucose) YPGal (+2% galactose), WOD (+2% glucose) and WOEG (+2% ethanol, 2% glycerol). The following chemicals were used to supplement solid YPGal plates: 10 mM nicotinamide, 10 mM nicotinic acid, and 10 μM or 50 μM MG132 (all from Sigma-Aldrich Corp., St Louis, MO, USA), and 10mM nicotinamide riboside (MedKoo Biosciences) at concentrations reported previously (77, 78).

### Preparation of Cell Extracts for Protein Aggregation Analysis

Cell extracts were prepared according to the protocol outlined in (Ocampo et al., 2013). Briefly, fifty-milliliter cultures (OD^600^ = 0.125) of strains expressing either polyQ-GFP or α-synuclein–GFP together or not with overexpression of the salvage NAD^+^ biosynthetic proteins Nma1, Npt1, Pnc1 or Qns1, were induced for 2, 4, 6, 8, and 24 h in complete media containing 2% galactose. Cells were harvested by centrifugation at 1100× *g* for 5 min and washed once with 1.2 M sorbitol. The cell wall was digested with 1 mg/ml zymolyase for 30 min, and the spheroplasts were re-suspended in lysis buffer (40 mm 4-(2-hydroxyethyl)-1-piperazineethanesulfonic acid buffer pH 7.4, 50 mm KCl, 1% Triton X-100, 0.3 mg/ml dithiothreitol, 5 mm EDTA and 0.5 mm phenylmethylsulfonyl fluoride). Samples were collected following gravity sedimentation of cell debris for 1 h on ice. The samples (T) were centrifuged 10 min at 800*g* at 4°C. The supernatant (S) was collected, and the pellet (P) was washed once in lysis buffer and then resuspended in the same volume of lysis buffer. Protein concentration was quantified by Folin method. Equal amount of protein was resuspended in Laemmli buffer [1% SDS, 0.1% (v/v) glycerol, 0.01% β-mercaptoethanol and 50 mm Tris–HCl pH 6.8) for immunoblot analysis.

### Recombinant Protein Purification

Yeast *NMA1, NPT1, PNC1*, and *QNS1* genes were cloned into the pET-41b(+) expression vector and then transformed into BL21 or BL21-star chemically competent *E. coli* cells (New England Biolabs, Ispwich, MA, USA). *E. coli* containing the expression plasmids were grown in complete LB broth (Thermo-Fisher Scientific, Waltham, MA, USA) at 30°C with shaking until cell cultures reached an OD 600 of approximately 0.5, at which point the temperature was lowered to 16°C and cultures were allowed to grow until reaching an OD 600 of approximately 0.7. Yeast recombinant protein expression was then induced by the addition of isopropyl β-D-1-thiogalactopyranoside (IPTG) (MilliporeSigma, Burlington, MA, USA) to the final concentration of 0.1mM, after which cultures were grown for an additional 12 to 72 hours at 16°C. Cells were collected and resuspended in a solution containing 50 mM sodium phosphate, 300 mM NaCl, yeast protease inhibitor cocktail (MilliporeSigma, Burlington, MA, USA), 1mM PMSF, 30 μg/ml benzamidine, and 10 μg/ml DNAse I and lysed at 12,000 psi using a French Press. The resulting lysate was clarified by centrifugation. Recombinant proteins were extracted from this crude lysate by affinity purification with TALON Affinity Resin (Takara Bio USA, Mountain View, CA, USA), washed with a solution containing 50 mM sodium phosphate, 300 mM NaCl, and 50 mM imidazole, and then eluted with a solution containing 50 mM sodium phosphate, 300 mM NaCl, and 200 mM imidazole. Buffer exchange was performed by running extracts through Sephadex resin columns (GE Healthcare, Chicago, IL, USA) and eluting with HEPES buffer (pH 7.4). Extracts were tested for purity by SDS page gel and Coomassie blue staining. The company GenScript purified a batch of Nma1 and Nma1Dct proteins for us.

### Citrate synthase aggregation assay

Citrate synthase (CS) aggregation measurements were performed as described previously (18). Substrate CS (Sigma, St. Louis, MO) was desalted and mixed with either egg white lysozyme (Sigma), purified catalytically active or inactive Nma1, Npt1, Pnc1 or Qns1 (purified in house), Nma1 and Nma1Δct (purified by GenScript (Piscataway, NJ) at varying concentrations (CS/protein: 1:0, 1:0.5, 1:1, 1:2, and 1:4) in HEPES (pH 7.4) buffer for 30 min. Aggregation of denatured CS was initiated at 43°C and was monitored as Raleigh scattering absorbance at 360 nm as a function of time. A FluoStar Optima plate reader (BMG Labtech, Cary, NC) was used for absorbance measurements. The relative chaperone activity of NMNAT was calculated as the scattering of CS aggregates with time versus NMNAT concentration.

### Luciferase Aggregation Assays

The luciferase aggregation assay was performed as described (31, 79), with a few modifications. HEK293T cells were transfected with pCMV-*cyt*Luciferase and one of the following plasmids: pCMV6-*HSP70*, pCMV6-*NMA1*, pCMV6-*NPT1*, pCMV6-*PNC1*, pCMV6-*QNS1,* pCMV6-*NMA1Δct*, empty pCMV6, using Lipofectamine (Invitrogen, Grand Island, NY). 48 h post-transfection, the cells were treated with the protein synthesis inhibitor cycloheximide (50 μg/mL for 30 min). Whereas one aliquot of cells was lysed immediately with luciferase lysis buffer (Promega, Madison, WI), two additional similar aliquots (second and third) were heat-shocked at 42°C for 15 min (which induced efficient unfolding of luciferase without killing the cells). The second aliquot of cells was lysed immediately after heat shock, and the third aliquot was allowed to recover at 37°C for 3 h. Luciferase activity was measured with the Luciferase Assay System (Promega, Madison, WI).

### Fluorescence microscopy

Wide-field fluorescence microscopy was performed to detect polyQ-GFP expression. We used an Olympus fluorescence BX61 microscope equipped with Nomarski differential interference contrast (DIC) optics, a Uplan Apo 100× objective (NA 1.35), a Roper CoolSnap HQ camera, and Sutter Lambda10-2 excitation and emission filter wheels, and a 175-watt Xenon remote source with liquid light guide. Images were acquired using SlideBook 4.01 (Intelligent Imaging Innovations, Denver, CO, USA). All experiments were performed at least in triplicate, with a minimum of 150 cells per sample.

### Immunoprecipitation assays

For Nma1-Hsc82 interaction analysis, cells expressing Nma1-HA were converted to spheroplasts by incubation in zymolyase buffer (1.2 M sorbitol, 0.4 mg/ml zymolyase, 20 mM K3PO4 pH 7.4) for 30 min at 30°C, pelleted by centrifugation at 6,000g for 10 min, and then solubilized in extraction buffer (50 mM Tris-HCl, pH 7.5, 100 mM KCl, 20 mM MgCl2, 1% digitonin, 10% glycerol, 1 mM PMSF, 1× EDTA-free protease inhibitor cocktail) and incubated for 30 min at 4°C. The lysates were cleared by centrifugation at 15,000 g for 15 min at 4°C. The lysate was incubated with 50 μl of washed recombinant protein A agarose beads (control) or anti-HA affinity gel beads at 4°C for 4 h with gentle rotation. The supernatant containing unbound material was subsequently collected, and the beads were washed four times with wash buffer (50 mM Tris-HCl, pH 7.5, 100 mM KCl, 20 mM MgCl2, 10% glycerol, 1 mM PMSF, 1× EDTA-protease inhibitor cocktail). The bound proteins were eluted using 1× Laemmli buffer containing 300 mM DTT and incubated at 95°C for 5 min. The different fractions were analyzed by immunoblotting using antibodies against HA, or Hsc82.

### Miscellaneous procedures

Standard procedures were used to prepare and ligate DNA fragments, transform and recover plasmid DNA from *E. coli,* and transform yeast. Western blots were treated with antibodies against the appropriate proteins followed by a second reaction with anti-mouse or anti-rabbit IgG conjugated to horseradish peroxidase (Sigma, St. Louis, MO, USA). The SuperSignal West Pico and Femto substrate kits (Pierce, Rockford, IL, USA) were used for the final detection.

### Statistical analysis

All experiments were performed at least in triplicate. Data are presented as means ± SD or SEM, as appropriate, of absolute values or percent of control. Values were compared by t-Student test or One-Way ANOVA, as noted in the figure legends. P < 0.05 was considered significant. Statistical analysis of citrate synthase aggregation assay measurements was performed with Graphpad Prism 6 software.

## Supplementary Material

Supplementary data includes 10 Figures.

### FUNDING

This work was supported by a Research Grant from The Army Research Office (ARO) # W911NF-16-1-0311 (to AB), a Merit Award from the Veterans Administration (VA) Biomedical Laboratory Research and Development 1I01BX003303-01 (to AB), and an HD Human Biology Project Fellowship from the Huntington’s Disease Society of America (HDSA) # GR009606 (to AR).

## ACKNOWLEDGEMENTS

We thank Yi Zhu and Dr. Grace Zhai for sharing reagents and protocols for CS aggregation studies and Julia Denissova for technical assistance. We thank Dr. Flavia Fontanesi for critical reading of the manuscript.

## AUTHORS CONTRIBUTIONS

AB designed the project; MP finished the experimental work, purified some recombinant proteins, obtained some data with the polyQ models and all data with the α-Syn models; AR started the experimental work, purified most recombinant proteins, and obtained data with the polyQ models; VB performed the protein aggregation assays and some microscopy studies; EN developed the luciferase assays; MP, AR, and AB wrote the manuscript.

## Authors emails

Meredith Pinkerton

Andrea Ruetenik

Viktoriia Bazylianska

Eva Nyvltova

Antoni Barrientos

## Data sharing plan

All the reagents will be distributed freely upon request.

## SUPPLEMENTAL FIGURES

**Figure S1.**
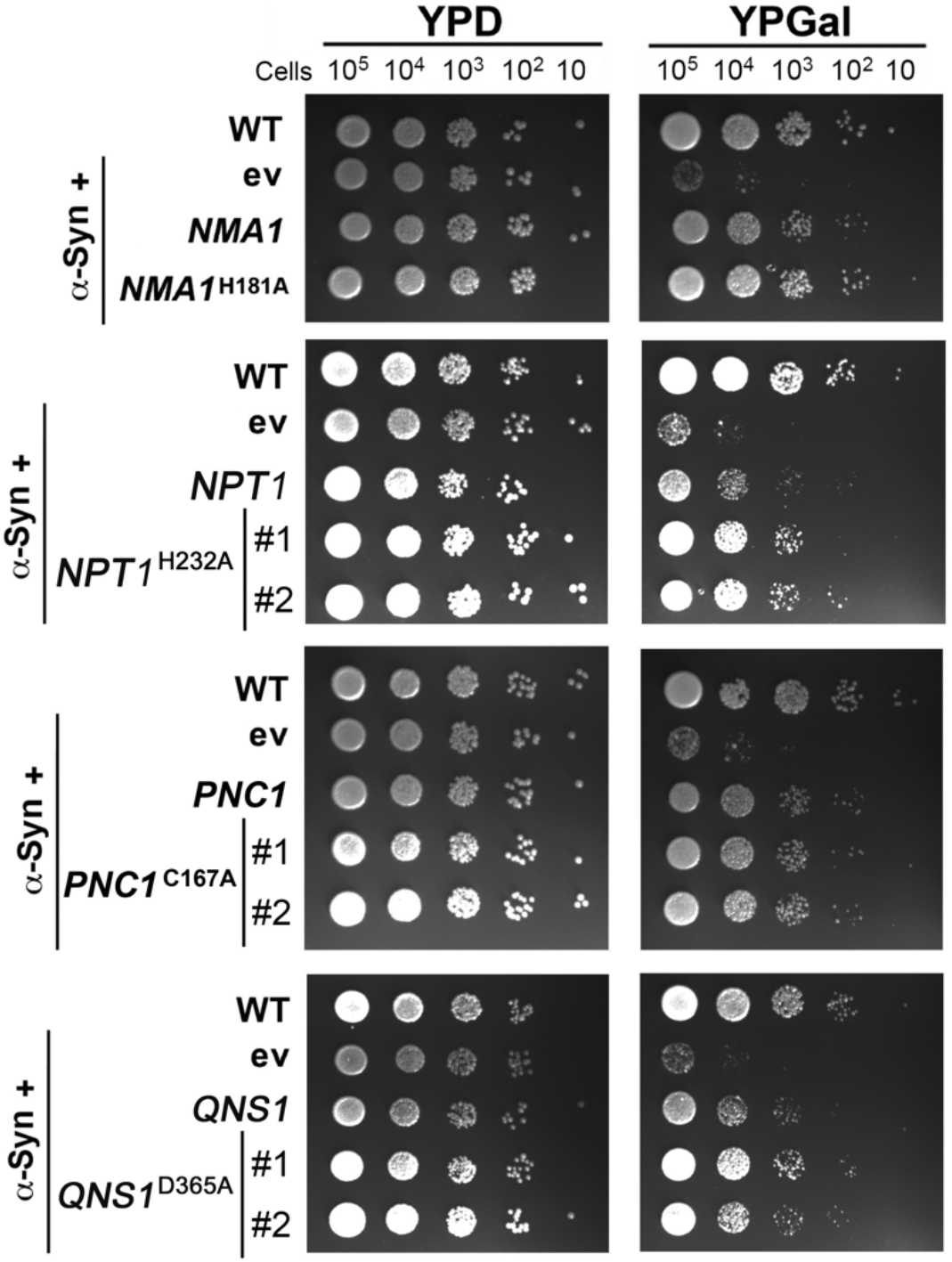
Catalytically inactive NAD^+^ salvage enzymes suppress α-synuclein-induced proteotoxicity in yeast. Serial dilutions growth test of wild-type cells (WT) expressing or not α-synuclein (α-Syn) and either an empty vector or catalytically active or inactive variants of the indicated salvage NAD^+^ pathway genes, under the control of a galactose inducible promoter, in solid non-inducing media (YPD) and inducing galactose media (YPGal). Where two independent clones were used, it is indicated by #. Pictures were taken after 2 days of growth.

**Fig S2.**
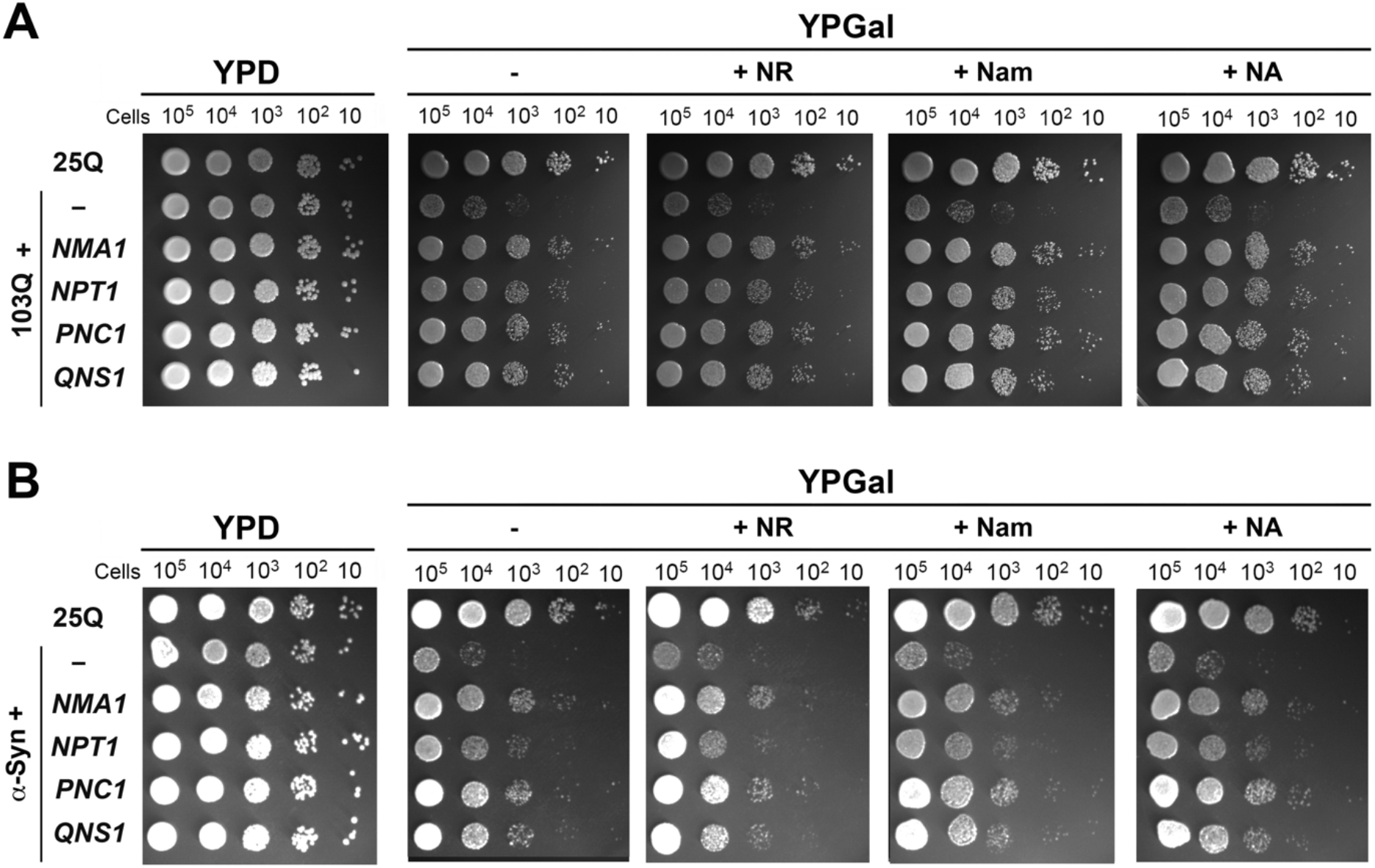
Boosting NAD^+^ biosynthesis with NAD^+^ precursors do not ameliorate 103Q- or α-Syn-induced proteotoxicity and is not additive to the protection exerted by overexpression of salvage NAD^+^ biosynthetic proteins. Serial dilutions growth test of wild-type cells expressing (**A**) non-toxic 25Q or toxic 103Q, and 103Q cells or (**B**) α-synuclein, expressing the indicated salvage NAD^+^ pathway genes, under the control of a galactose inducible promoter, in solid non-inducing media (YPD) and inducing galactose media (YPGal) supplemented with 10 mM of nicotinamide riboside (NR), nicotinamide (Nam) or nicotinic acid (NA). Pictures were taken after 2 days of growth.

**Fig S3.**
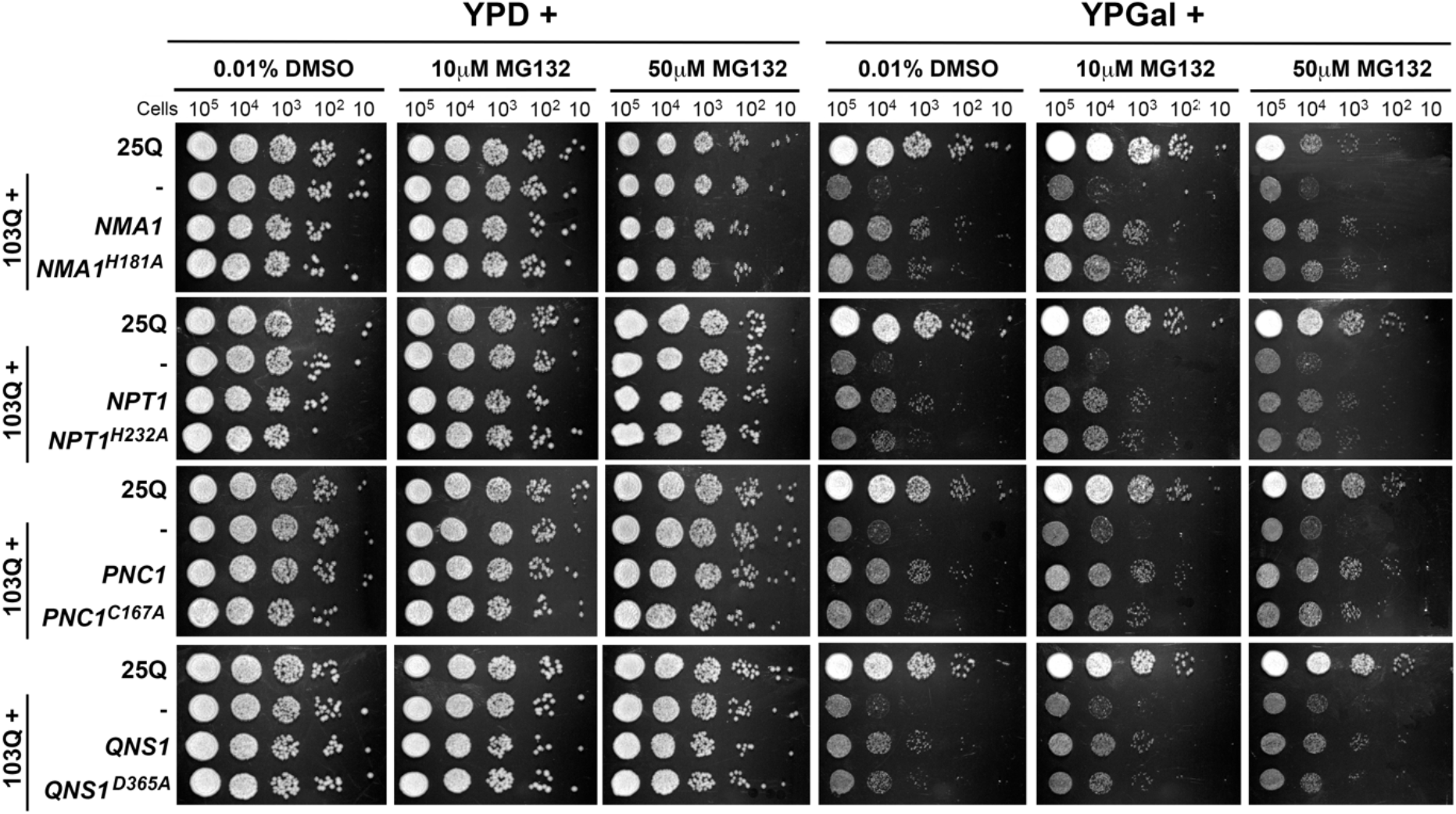
Inhibition of the proteasome with MG132 does not prevent the suppression of 103Q-induced toxicity by catalytically active or inactive NAD^+^ salvage enzymes. Serial dilutions growth test of wild-type cells expressing non-toxic 25Q or toxic 103Q, and 103Q cells expressing WT or catalytically inactive variants of the indicated salvage NAD^+^ pathway genes, under the control of a galactose inducible promoter, in solid non-inducing media (YPD) and inducing galactose media (YPGal) supplemented with 10 μM or 50 μM MG132 or the vehicle DMSO. Pictures were taken after 2 days of growth.

**Fig S4.**
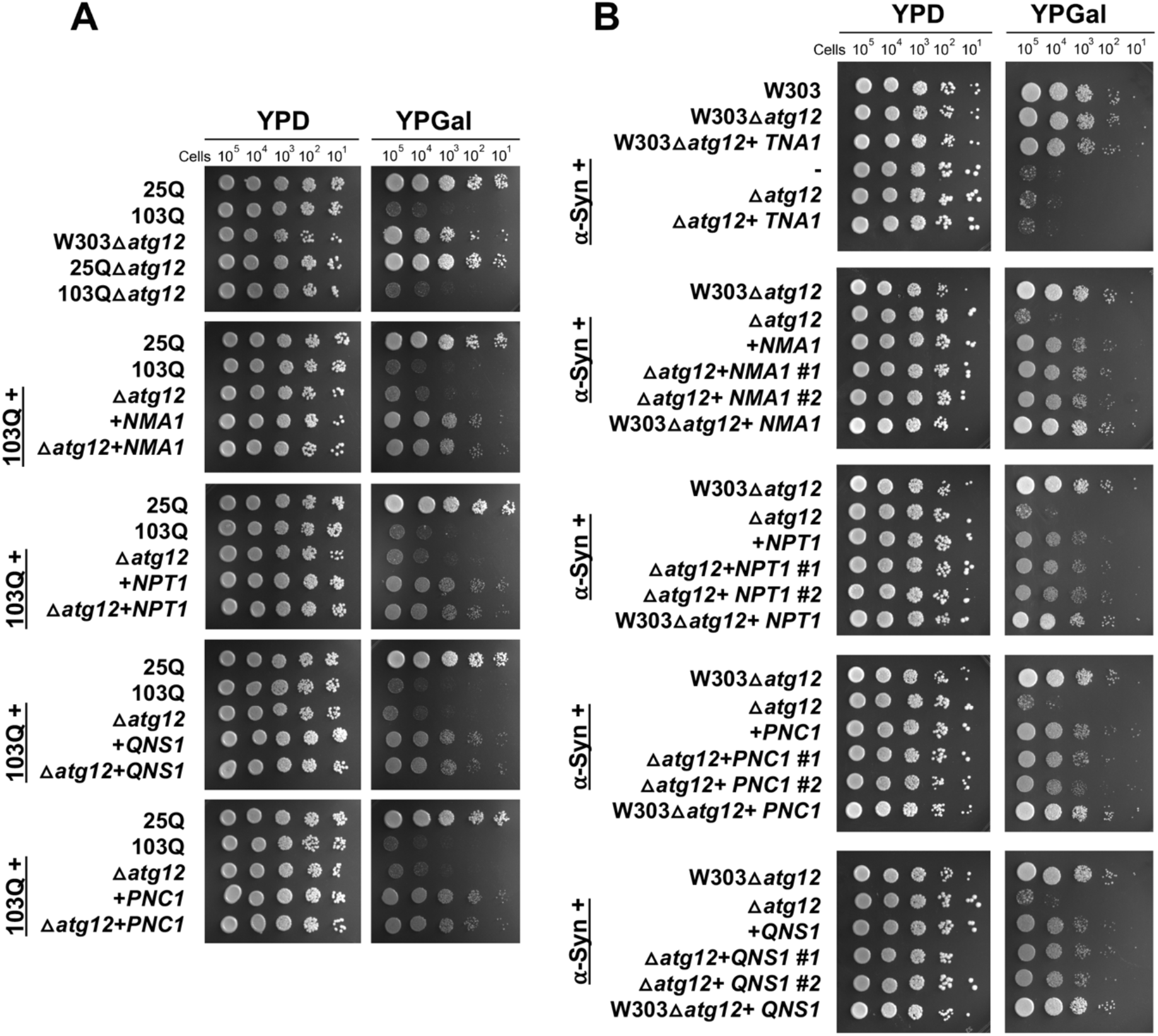
Deletion of *atg12*, essential for autophagosome formation, does not prevent the suppression of 103Q-induced toxicity by catalytically active or inactive NAD^+^ salvage enzymes. Serial dilutions growth test of wild-type cells expressing non-toxic 25Q or toxic 103Q, and 103Q cells deleted or not for *atg12*, expressing WT or catalytically inactive variants of the indicated salvage NAD^+^ pathway genes, under the control of a galactose inducible promoter, in solid non-inducing media (YPD) and inducing galactose media (YPGal). Pictures were taken after 2 days of growth.

**Fig S5.**
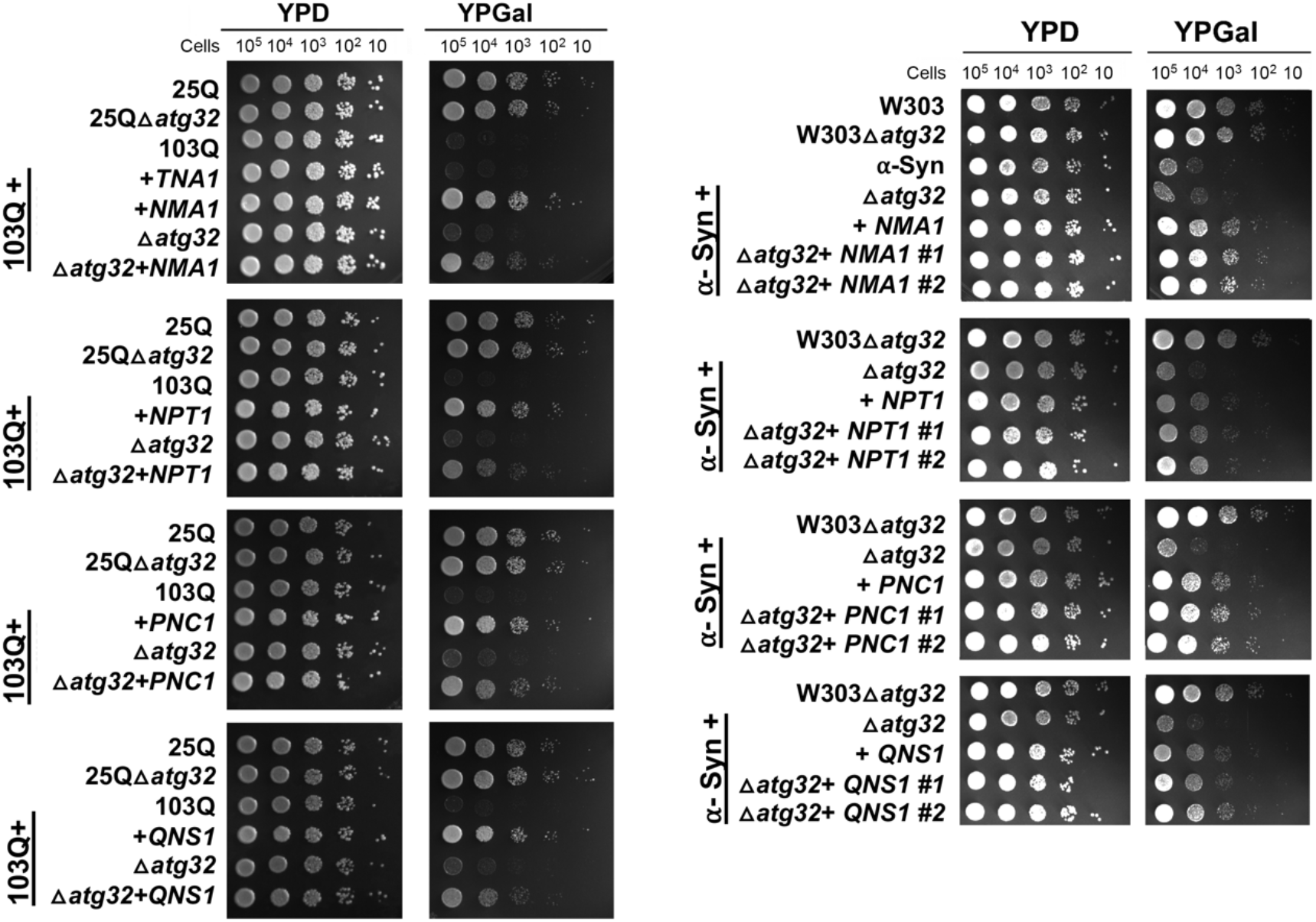
Deletion of *atg32*, required to initiate mitophagy, does not prevent the suppression of 103Q-induced toxicity by catalytically active or inactive NAD^+^ salvage enzymes. Serial dilutions growth test of wild-type cells expressing non-toxic 25Q or toxic 103Q, and 103Q cells deleted or not for *atg12*, expressing WT or catalytically inactive variants of the indicated salvage NAD^+^ pathway genes, under the control of a galactose inducible promoter, in solid non-inducing media (YPD) and inducing galactose media (YPGal). Pictures were taken after 2 days of growth.

**Fig S6.**
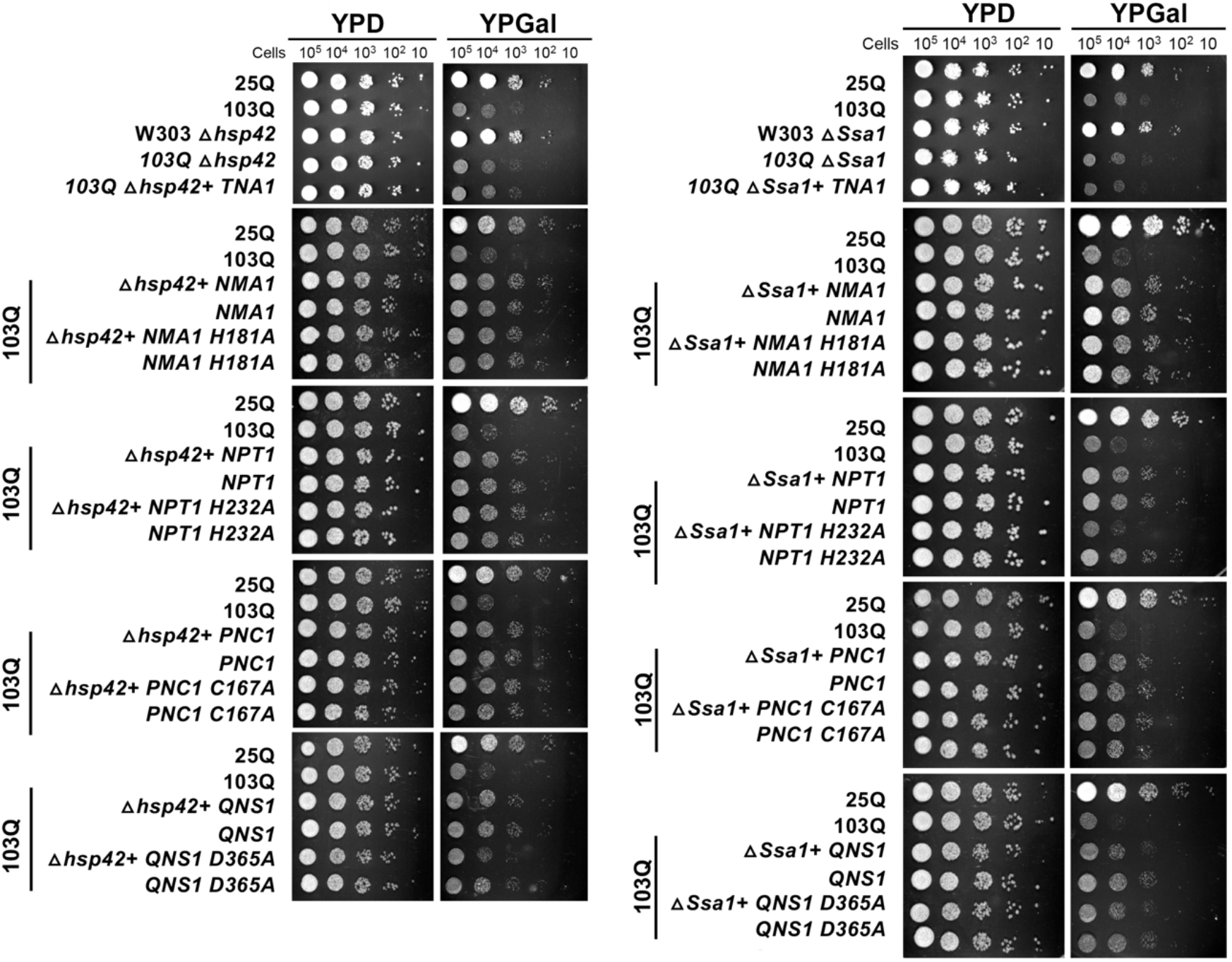
Deletion of the heat-shock proteins Hsp42 or Hsp70 (Ssa1), does not prevent the suppression of 103Q-induced toxicity by catalytically active or inactive NAD^+^ salvage enzymes. Serial dilutions growth test of wild-type cells expressing non-toxic 25Q or toxic 103Q, and 103Q cells deleted or not for *hsp42* or *ssa1*, expressing WT or catalytically inactive variants of the indicated salvage NAD^+^ pathway genes, under the control of a galactose inducible promoter, in solid non-inducing media (YPD) and inducing galactose media (YPGal). Pictures were taken after 2 days of growth.

**Fig S7.**
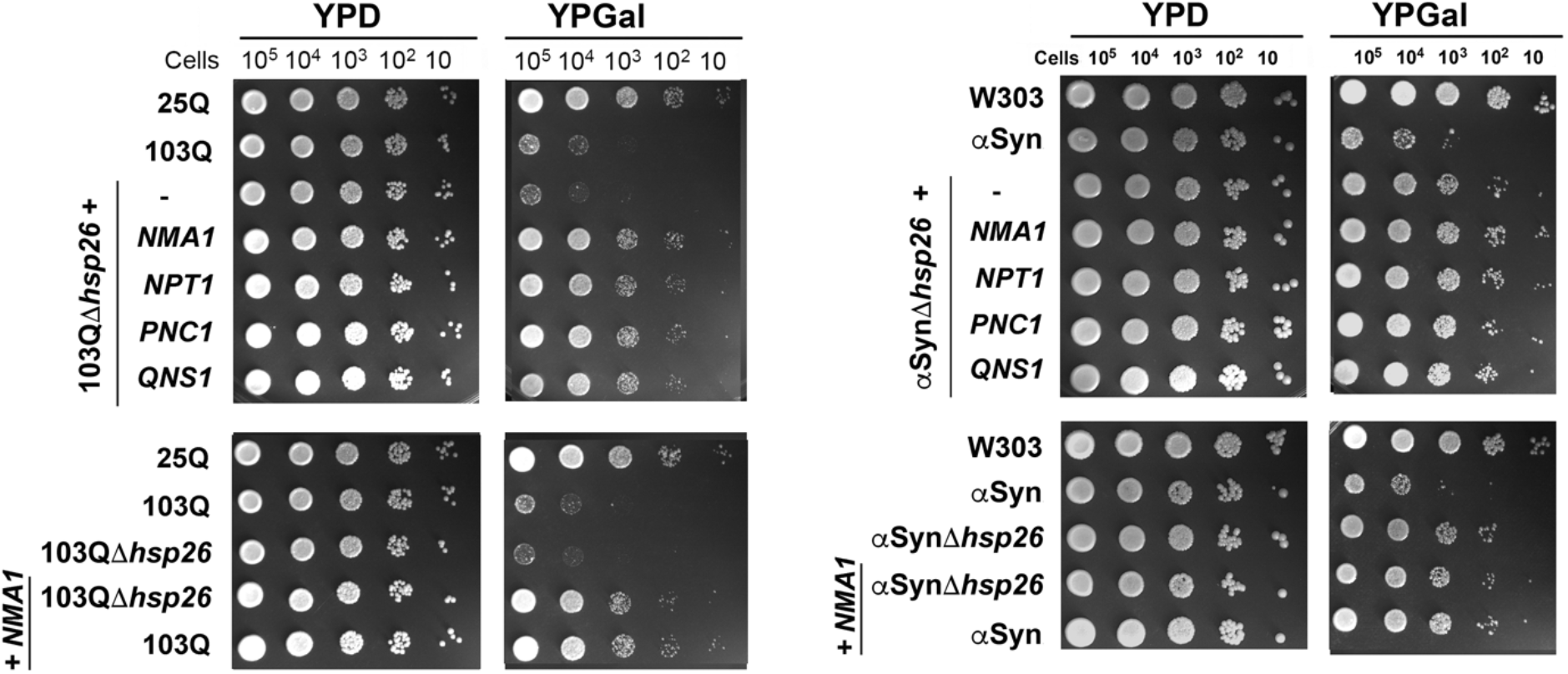
Deletion of the heat-shock protein Hsp26, does not prevent the suppression of 103Q- and α-Syn-induced toxicity by catalytically active or inactive NAD^+^ salvage enzymes. Serial dilutions growth test of wild-type cells expressing toxic α-Syn, non-toxic 25Q or toxic 103Q, and 103Q cells deleted or not for *hsp26*, expressing WT or catalytically inactive variants of the indicated salvage NAD^+^ pathway genes, under the control of a galactose inducible promoter, in solid non-inducing media (YPD) and inducing galactose media (YPGal). Pictures were taken after 2 days of growth.

**Fig S8.**
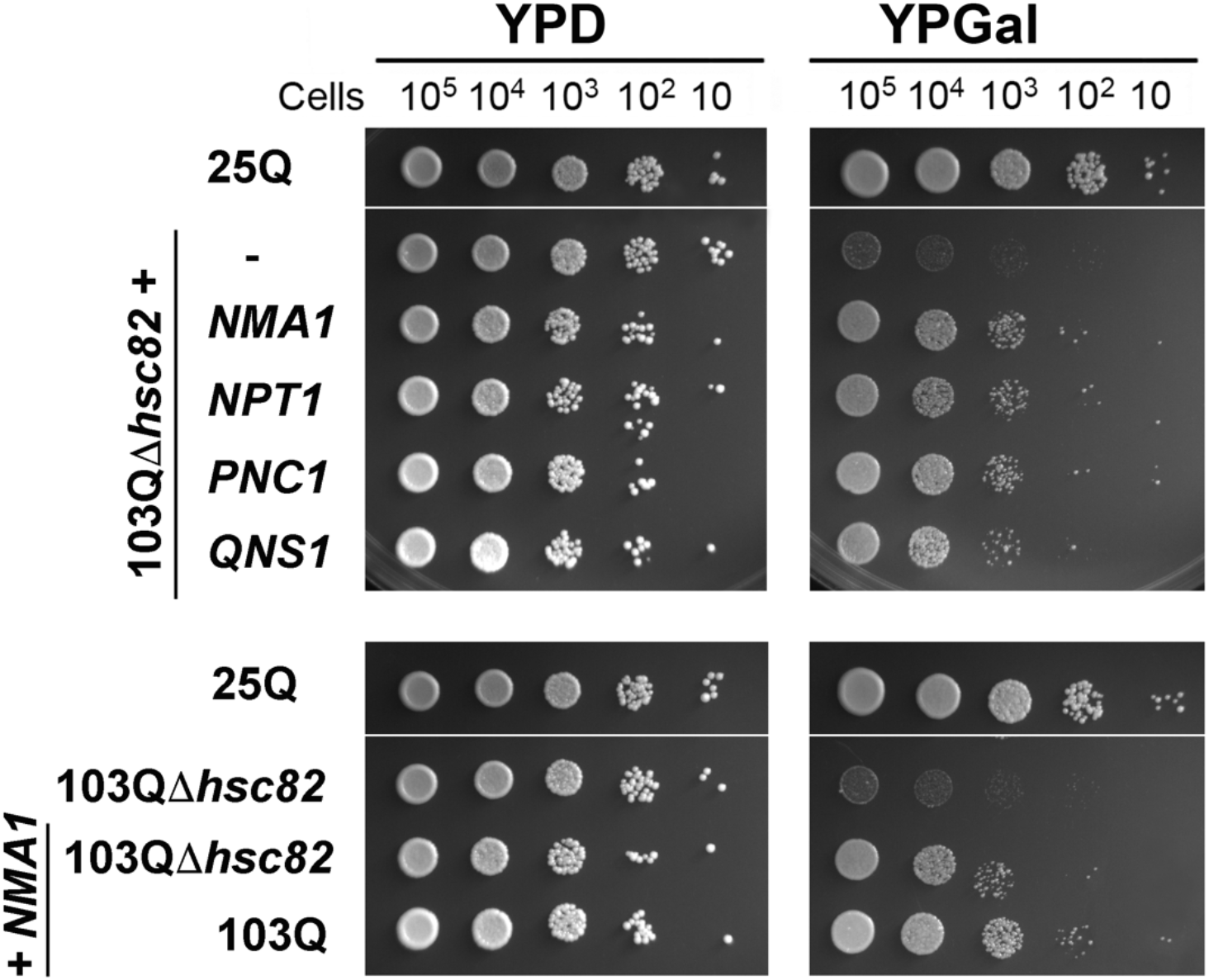
Deletion of the heat-shock protein Hsc82, does not prevent the suppression of 103Q-induced toxicity by catalytically active or inactive NAD^+^ salvage enzymes. Serial dilutions growth test of wild-type cells expressing, non-toxic 25Q or toxic 103Q, and 103Q cells deleted or not for *hsc82* expressing WT or catalytically inactive variants of the indicated salvage NAD^+^ pathway genes, under the control of a galactose inducible promoter, in solid non-inducing media (YPD) and inducing galactose media (YPGal). Pictures were taken after 2 days of growth.

**Fig S9.**
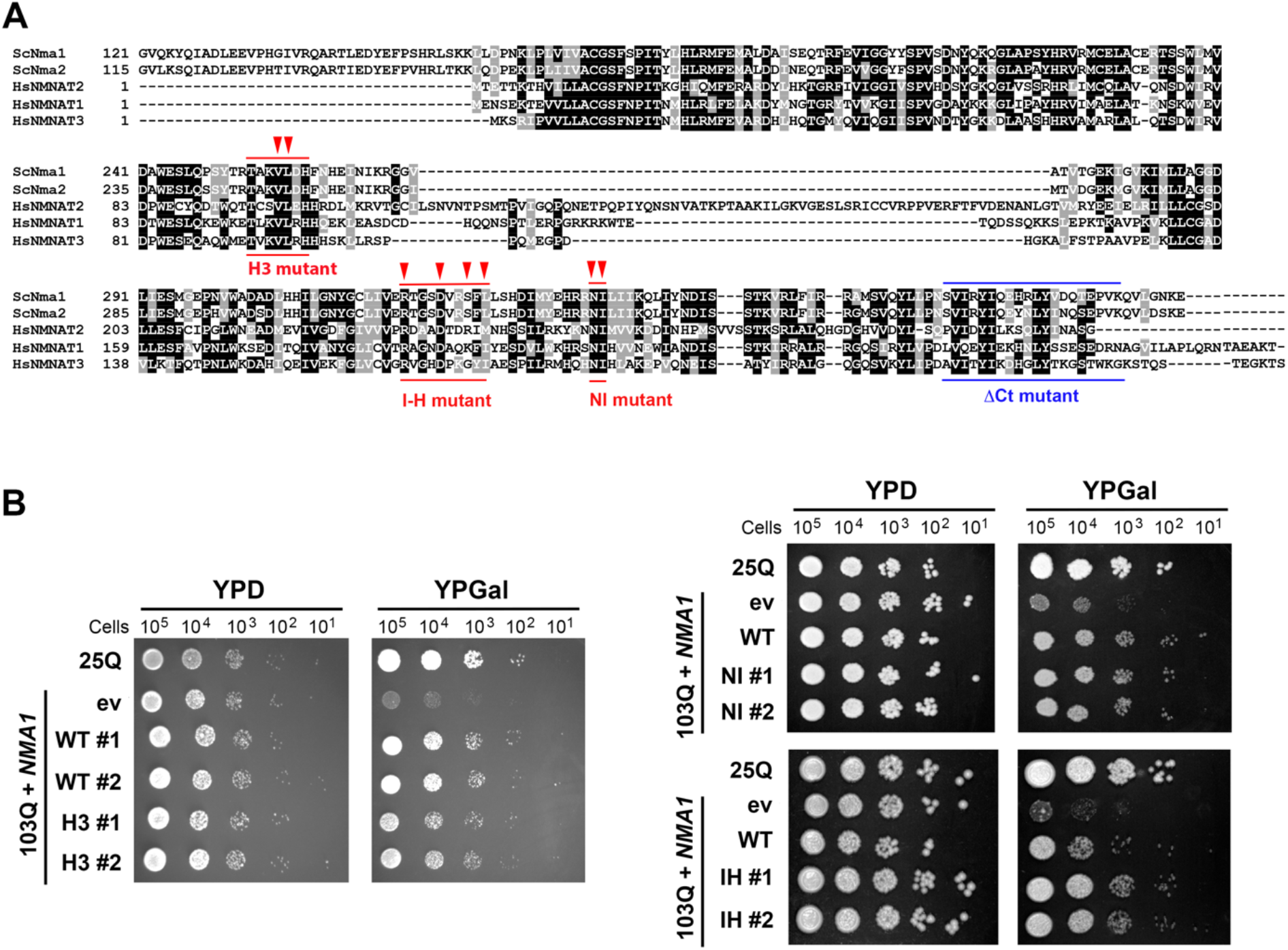
Mutagenesis of key residues in Nma1 does not affect its ability to suppress 103Q- and α-Syn-induced proteotoxicity. (**A**) Multiple protein alignment of *Saccharomyces cerevisiae* Nma1 and Nma2 and *Homo sapiens* NMNAT1, NMNAT2, and NMNAT3 using the Clustal Omega program from EMBL-EBI. The mutagenized domains and residues are highlighted in red. These are: (i) helix 3 (H3), conserved in the bacterial UspA chaperone, which includes Nma1 amino acids 252-TAKVLDH-259 (1), in which VL were changed to EE; (ii) helix 4 which includes Nma1 amino acids 220-RTGSDVRSFL-229 and occupies a internal position in the hexameric NMNAT1 structure (I-H), in which the sequence was changed to 220-ATGSAVRSFA-229; and (iii) two conserved residues, Nma1 341-NI-342 (NI), located in a hotspot region for human NMNAT1 mutations associated with Leber congenital amaurosis 9 (LCA9) who retain a substantial catalytic activity (2). The C-terminal fragment deleted in the *NMA1Δ*ct yeast strain is marked in blue. (**B**) Serial dilutions growth test of wild-type cells expressing non-toxic 25Q or toxic 103Q, and 103Q cells expressing WT or the Nma1 variants H3, I-H or NI described in panel (A), under the control of a galactose inducible promoter, in solid non-inducing media (YPD) and inducing galactose media (YPGal). Pictures were taken after 2 days of growth.

**Fig. S10.**
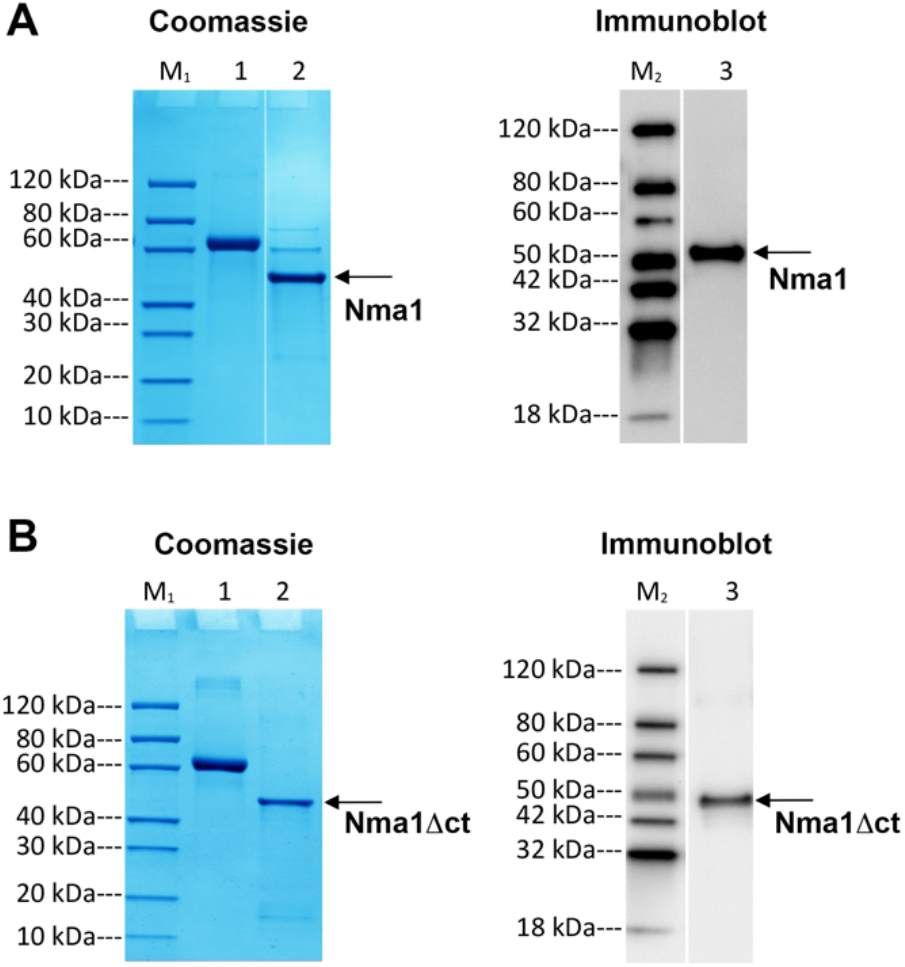
Analysis of recombinant full-length Nma1 and truncated Nma1Δct proteins expressed and purified from *E. coli*. Coomassie staining and immunoblot analyses of (**A**) Nma1 and (**B**) Nma1Δct purified recombinant proteins separated SDS-PAGE. Lane M1: Protein Marker, GenScript, Cat. No. M00516 Lane M2: Protein Marker, GenScript, Cat. No. M00521 Lane 1: BSA (2.00 μg) Lane 2: Purified Nma1 variant (Reducing condition, 2.00 μg) Lane 3: Purified Nma1 variant (Reducing condition). Primary antibody: Mouse-anti-His mAb (GenScript, Cat.No. A00186)

**Fig S11.**
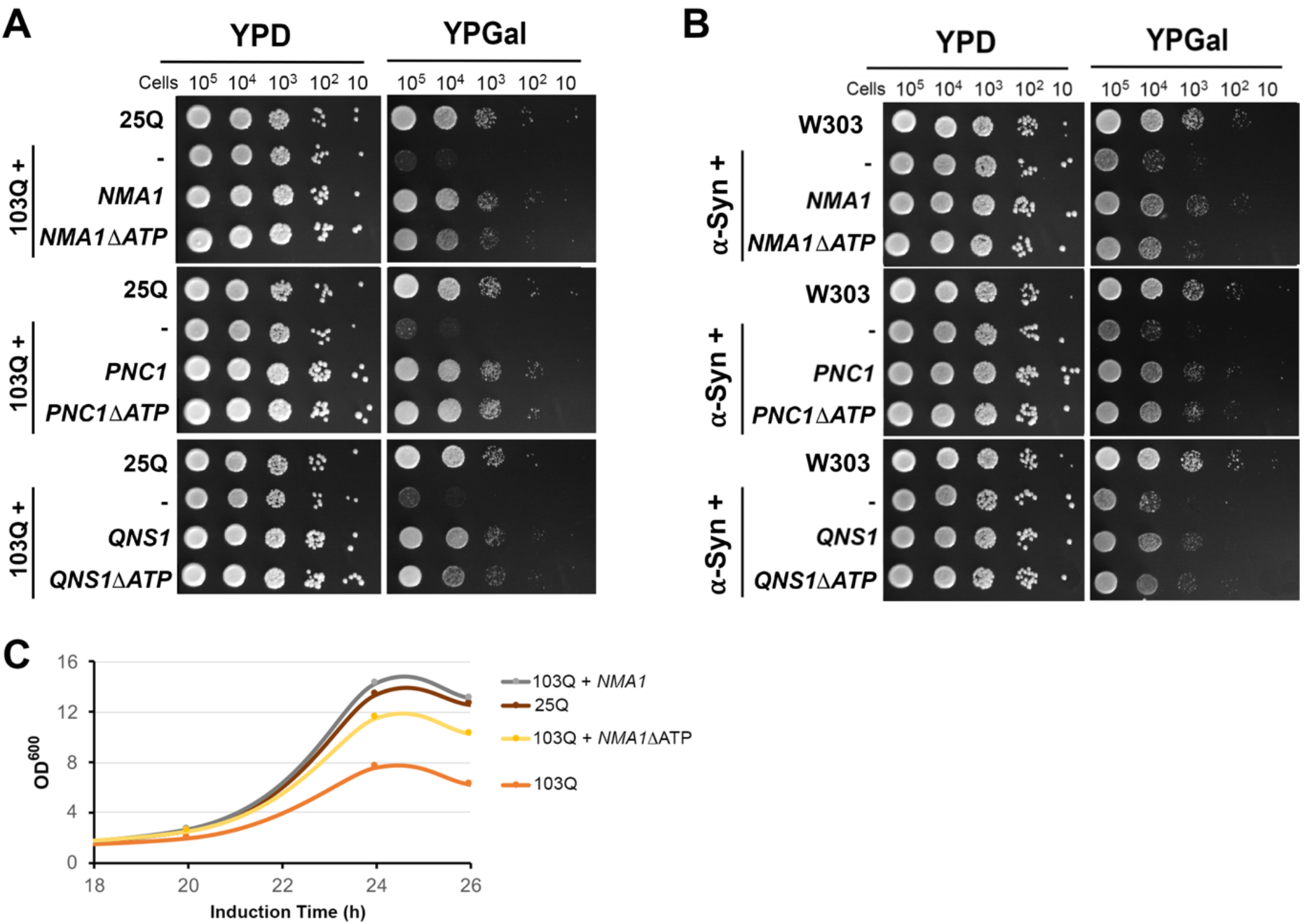
Mutagenesis of ATP-binding motifs in Nma1 but no other salvage NAD^+^ biosynthetic proteins slightly affect its ability to suppress 103Q- and α-Syn-induced proteotoxicity. (**A**) Serial dilutions growth test of wild-type cells expressing non-toxic 25Q or toxic 103Q, and 103Q cells expressing WT or the Δ ATP variants of NMA1, PNC1, or QNS1, under the control of a galactose inducible promoter, in solid non-inducing media (YPD) and inducing galactose media (YPGal). Pictures were taken after 2 days of growth. (**B**) Growth curve in liquid synthetic media containing galactose of the indicated strains. Measurements of absorbance at 600nm were performed up to 26 h after induction.

